# Three-hinge gyri link individualized folding topology to cortical and superficial white-matter diffusion organization

**DOI:** 10.64898/2026.10.14.757725

**Authors:** Minheng Chen, Xiuyu Dong, Jiale Cheng, Dan Hu, Tianming Liu, Dajiang Zhu, Gang Li

## Abstract

Individual differences in cortical folding are a hallmark of human brain anatomy, but what this variability reflects in the underlying tissue remains unclear. Three-hinge gyri (3HGs), where three gyral crests converge, provide an individualized higher-order folding landmark for probing this relationship. Using 7T diffusion MRI in six postmortem hemispheres, we found lower mean, axial, and radial diffusivity and higher fractional anisotropy at 3HGs than at matched gyral crowns, with the principal tensor pattern replicated in 414 age-matched *in vivo* adults. Diffusivity differences extended across the cortical ribbon, whereas anisotropy- and fiber-related differences strengthened toward and beneath the white surface. *Ex vivo*, diffusion measures also varied with gyral-skeleton degree, and streamlines sampled beneath 3HGs preferentially reached other 3HGs. These findings link individualized folding topology to cortical and superficial white-matter diffusion-inferred tissue properties, providing tissue-level context for understanding the microstructural correlates of individual folding variation.

## 1 Introduction

Cortical folding is one of the most individually distinctive features of human brain anatomy. Although major gyri and sulci follow broadly conserved anatomical patterns, their detailed arrangement, depth, branching, and interruption vary substantially across individuals [1–3]. Increasing evidence suggests that this variability is behaviorally and clinically meaningful rather than merely geometric. Individual sulcal patterns have been associated with executive function, reasoning, language, and memory, while atypical folding has been reported in neurodevelopmental and psychiatric disorders such as autism and schizophrenia [4–9]. Many qualitative folding patterns are established early and remain stable across later development [10, 11], making them useful anatomical phenotypes for studying individual differences in brain organization. Yet what these individualized folding patterns reflect in the underlying cortical tissue remains poorly understood.

At the broader scale of gyri and sulci, studies indicate that surface morphology is associated with differences in tissue structure [12, 13]. Gyral crowns and sulcal fundi differ in cortical thickness, cellular and neuritic architecture, and the geometry of fibers approaching the cortex [14–16]. Whether such relationships extend to finer, individually variable folding topology is less clear. Three-hinge gyri (3HGs), where three gyral crests converge, provide a particularly informative landmark for addressing this question. Their number, location, and morphology vary markedly across individuals while retaining reproducible regional organization [1, 2]. Previous *in vivo* studies have associated 3HGs with greater cortical thickness, stronger structural connectivity, and broader functional involvement [17–22]. 3HGs also occupy communication-intensive positions within the structural network [22], and more recent work has identified reproducible communities within the gyral folding skeleton [23, 24]. Individualized folding connectomes have further been applied to characterize disease-related alterations [25–27]. These observations suggest that 3HGs are more than geometric junctions, but the tissue organization associated with these individualized folding landmarks has not been directly characterized. Several questions therefore remain. First, whether 3HGs exhibit a distinct cortical diffusion phenotype relative to ordinary gyral crowns, and whether such differences persist within the same cortical regions and after accounting for geometric and partial-volume effects, is unknown. Second, resolving their distribution across cortical depth and into superficial white matter is necessary to determine whether the phenotype is confined to the gray–white matter boundary or extends more broadly through the cortex. Third, comparing junctions with different branching degrees can establish whether any observed differences are specific to 3HGs or form part of a broader relationship between gyral convergence and tissue organization. Postmortem *ex vivo* MRI is particularly well suited to these questions. Prolonged, motion-free acquisitions provide submillimeter diffusion data, allowing the cortical ribbon and superficial white matter to be resolved directly. High-resolution structural images further enable specimen-specific reconstruction of the white and pial surfaces, so diffusion measures can be referenced to cortical geometry, sampled across cortical depth, and followed through the gray–white matter transition [28–32].

Here, we combined ultra-high-resolution *ex vivo* 7T diffusion MRI in six postmortem human hemispheres with independent *in vivo* validation in 414 age-matched adults. We found that 3HGs showed a reproducible low-diffusivity cortical phenotype relative to ordinary gyral crowns, with lower MD, AD, and RD consistently reproduced in the living cortex. The *ex vivo* data further showed that diffusivity differences extended across the cortical ribbon, whereas anisotropy- and fiber-related differences became more pronounced toward the gray– white matter boundary and superficial white matter. These diffusion properties also varied systematically with gyral-skeleton degree, placing 3HGs within a broader relationship between folding convergence and local tissue organization. Tractography provided complementary evidence of preferential inter-hinge wiring beneath these junctions. These findings indicate that individualized folding patterns are not only differences in surface geometry, but are also associated with systematic variation in cortical and superficial white-matter diffusion organization. This provides an anatomical context for interpreting folding phenotypes reported in cognitive and clinical studies.

## 2 Results

### 2.1 Three-hinge gyri exhibit a reproducible cortical diffusion signature

3HGs were distributed throughout the cortical folding skeleton and were more densely represented outside primary cortical regions in both datasets (Supplementary Fig. S5). They were contrasted with ordinary gyral-crown locations and sulcal fundi (Fig. 1a). To characterize their local tissue phenotype, we quantified cortical thickness together with diffusion measures within 2-mm geodesic neighborhoods centered on each landmark. Nearby 3HG detections whose sampling supports overlapped were aggregated into individual gyral junctions to avoid repeated measurement of the same cortical tissue. Across the six specimens, 3HGs showed a consistent diffusion profile relative to gyral crowns (Fig. 1b). Mean diffusivity (MD), axial diffusivity (AD), radial diffusivity (RD), Diffusion basis spectrum imaging (DBSI) hindered fraction, and DBSI free-water fraction were lower at 3HGs in all six specimens, whereas fractional anisotropy (FA) was consistently higher. Mean effect sizes ranged from *d* = −0.211 for RD to *d* = 0.171 for FA, with all six measures showing the same direction across specimens (*P* ≤ 0.010). DBSI fiber and restricted fractions showed smaller effects, whereas the most consistent differences were observed in tensor diffusivity, FA, and the hindered and free-water components. Cortical thickness was greater on average at 3HGs, but this effect was present in five of six specimens rather than uniformly across the cohort. Comparison with sulcal fundi placed the 3HG–crown differences within the broader gyral–sulcal axis (Fig. 1c), while subsequent within-region analyses tested whether these effects persisted independently of regional localization. The largest and most reproducible differences were therefore concentrated in diffusion-derived measures. These findings identify 3HGs as cortical landmarks with a reproducible low-diffusivity and more anisotropic diffusion phenotype. The spatial distribution of these effects was examined within individual Desikan–Killiany (DK) regions (Fig. 1d). Within-region comparisons preserved the direction of the principal diffusion effects across much of the cortex: MD and DBSI hindered fraction were lower at 3HGs in most cortical parcels, whereas regional variation was greater for thickness and fiber-related measures. Thus, the whole-cortex differences were not attributable solely to the preferential localization of 3HGs within a small number of cortical regions.

**Fig. 1.**
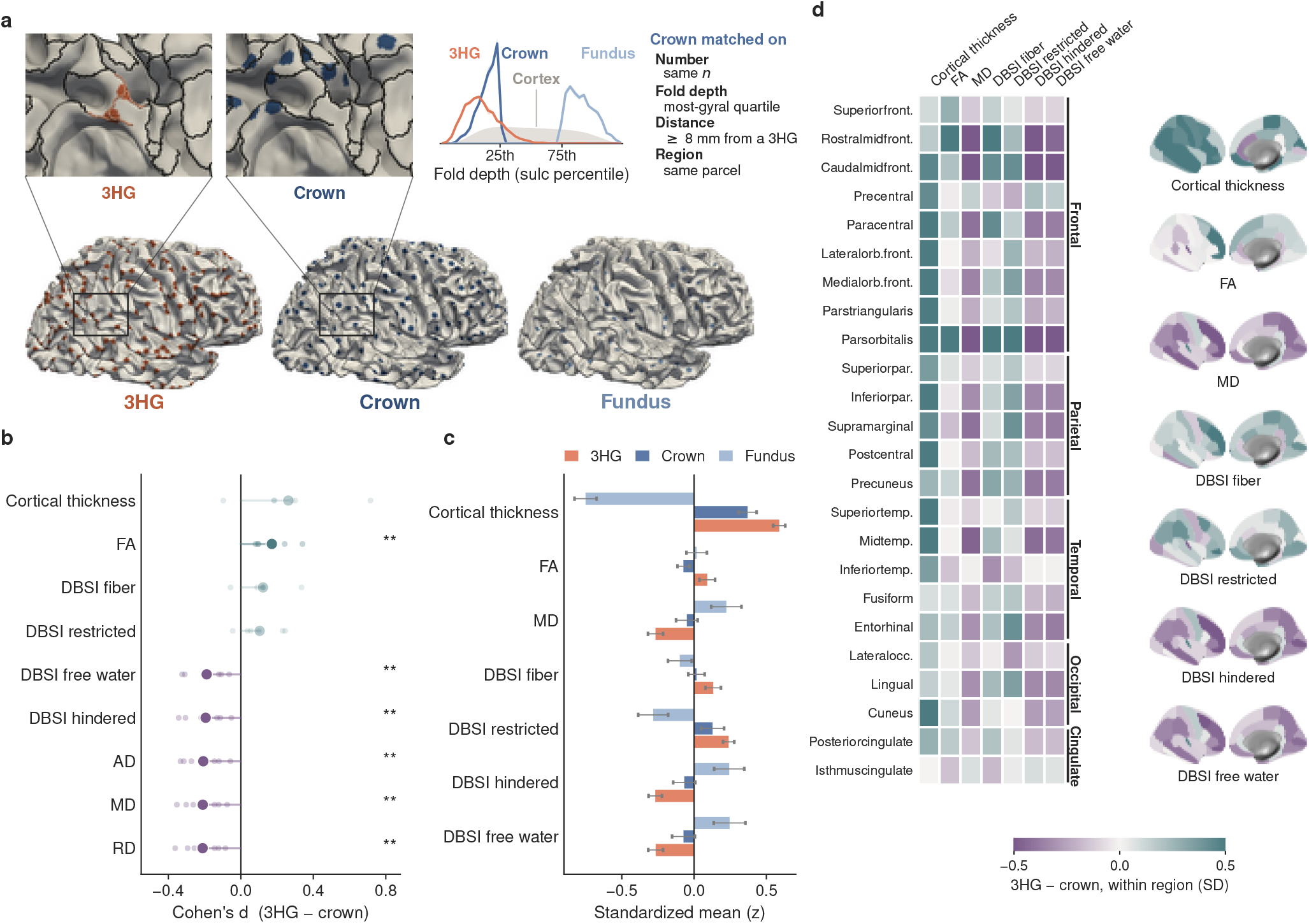
Three-hinge gyri show a distinct cortical diffusion phenotype. **a** Definition of the cortical sampling locations used throughout the analysis. Three-hinge gyri (3HGs, orange) were identified at junctions where three gyral crests converge. Ordinary gyral-crown controls (dark blue) were selected from highly gyral cortex and matched to 3HGs in number and cortical parcel, while being located at least 8 mm from the nearest 3HG. Sulcal-fundus locations (light blue) were included as a reference representing the opposite end of the folding profile. The fold-depth distributions illustrate the relative positions of 3HG, crown, and fundus samples along the gyral-to-sulcal dimension, with 3HGs and matched crowns occupying the gyral end of the distribution. Surface renderings show the spatial distribution of the three sampling classes on a representative hemisphere. **b** Differences between 3HGs and matched gyral crowns across six *ex vivo* hemispheres. Effect sizes are shown as Cohen’s *d* for cortical thickness and diffusion-derived measures. Small circles indicate individual hemispheres and large circles indicate the across-hemisphere mean. Negative values indicate lower measurements at 3HGs and positive values indicate higher measurements. Color hue indicates the predominant direction of the 3HG–crown difference, while color intensity reflects the consistency of that direction across hemispheres. **c** Standardized values at 3HG, crown, and fundus locations for selected cortical measures, illustrating how the 3HG phenotype relates to the broader gyral-to-sulcal profile. Bars show mean standardized values with error bars across hemispheres. **d** Regional consistency of the 3HG–crown phenotype across Desikan–Killiany cortical parcels. The heat map shows the within-region standardized difference between 3HGs and matched crowns for each measure, expressed as 3HG minus crown in standard-deviation units. Cortical surface maps show the spatial distribution of these regional effects. The sign and magnitude of each effect are indicated by the color scale. AD, axial diffusivity; FA, fractional anisotropy; MD, mean diffusivity; RD, radial diffusivity; DBSI, diffusion basis spectrum imaging. AD and RD were omitted from **c** and **d** because their regional patterns closely paralleled MD, but both were included in the full statistical analysis shown in **b**. P *<* 0.01, two-sided one-sample t-test against zero.

### 2.2 The 3HG diffusion phenotype extends across cortical depth and into superficial white matter

We next asked whether the 3HG-associated diffusion phenotype reflected a localized boundary effect or extended through the cortical ribbon. Nine equivolumetric surfaces were generated between the white and pial boundaries to account for curvature-dependent variation in cortical layer geometry (Fig. 2a). Statistical comparisons were performed within DK parcels, such that 3HG samples were compared with gyral-crown samples within the same DK parcel. MD, DBSI hindered fraction, and DBSI free-water fraction showed a robust main effect of landmark class across cortical depth (Fig. 2d). All three measures were consistently lower at 3HGs across the six specimens, with FDR-corrected *q* = 0.032. Their 3HG–crown differences changed little across the cortical ribbon, and no significant depth interaction was detected. These results indicate that the lower-diffusivity phenotype of 3HGs is not confined to the immediate gray–white matter boundary, but extends across intracortical depth. A different pattern was observed for measures related to diffusion anisotropy. DBSI fiber fraction and FA did not show a significant average 3HG–crown offset across the entire cortical ribbon, but their differences varied systematically with cortical depth, with significant class-by-depth interactions after correction for multiple comparisons (*q* = 0.026 for both). DBSI restricted fraction showed a similar but weaker trend (*q* = 0.056). Thus, the intracortical organization at 3HGs separates into two components: a depth-consistent reduction in diffusivity-related measures and a depth-dependent modulation of anisotropic/fiber-related diffusion.

**Fig. 2.**
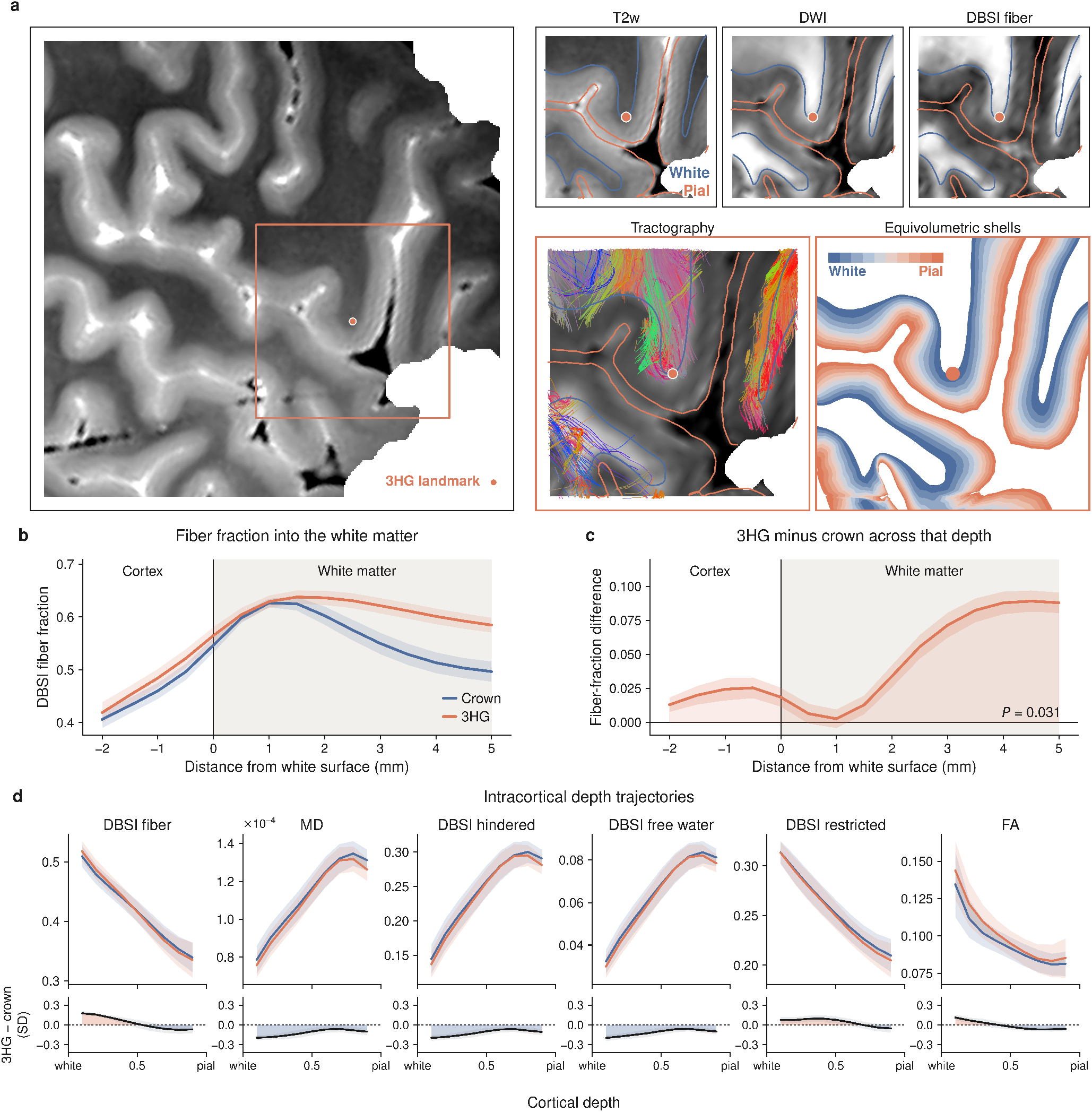
Depth-resolved diffusion differences at three-hinge gyri extend from the cortex into superficial white matter. **a** Depth-resolved sampling around a representative three-hinge gyrus (3HG; orange dot). The overview T2-weighted image shows the local cortical geometry and the region enlarged in the adjacent panels. Enlarged views show the T2-weighted image, diffusion-weighted image, DBSI fiber fraction, and local tractography with the reconstructed white and pial surfaces. Nine equivolumetric surfaces spanning the white-to-pial cortical ribbon were used for intracortical sampling. **b** DBSI fiber-fraction profiles sampled along the local surface normal across the gray–white matter boundary at 3HG and matched crown locations. Distance is referenced to the white surface at 0 mm, with negative values extending into the cortex and positive values extending into superficial white matter. The 3HG and crown profiles are similar within the cortex but progressively separate beneath the white surface, with higher DBSI fiber fraction beneath 3HGs. **c** Corresponding 3HG–crown difference in DBSI fiber fraction along the same cortical-to-white-matter trajectory. Positive values indicate higher fiber fraction at 3HGs. The difference increases within superficial white matter and is positive across all six ex vivo hemispheres in the superficial white-matter segment (*P* = 0.031). **d** Intracortical profiles of DBSI fiber fraction, mean diffusivity (MD), DBSI hindered fraction, free-water fraction, restricted fraction, and fractional anisotropy (FA) across the nine equivolumetric surfaces from the white to pial boundary. Upper panels show the measurements at 3HG and matched crown locations. Lower panels show the corresponding within-region standardized 3HG–crown differences. Diffusivity-related differences remain relatively stable across cortical depth, whereas anisotropy- and fiber-related differences vary with depth and become more pronounced toward the gray–white matter boundary. DBSI, diffusion basis spectrum imaging.

Importantly, the depth-resolved diffusion effects were not attributable to tissue partial volume (Supplementary Fig. S1a). Gray-matter fractions were nearly identical between 3HG and crown samples (0.945 versus 0.945), with 96% of samples arising from voxels containing at least 70% gray matter. Restricting the analysis to these high-purity voxels left the effects essentially unchanged, whereas explicit adjustment for white-matter and non-brain fractions slightly increased the MD, hindered, and free-water differences, indicating that partial volume attenuated rather than generated the observed intracortical phenotype. The depth-dependent intracortical pattern prompted us to examine the transition from cortex into the underlying white matter. DBSI fiber fraction was sampled continuously from 2 mm above to 5 mm below the white surface (Fig. 2b,c). Within the cortical ribbon, 3HG and crown profiles were similar. After crossing the white surface, however, the two profiles progressively diverged. The DBSI fiber fraction remained higher beneath 3HGs, with the difference increasing toward deeper sampled white matter and reaching approximately 0.08 at 4-5 mm below the white surface. This subcortical difference was positive in all six specimens (*P* = 0.031). This pattern suggests that the fiber-related component of the 3HG phenotype is spatially concentrated toward the cortical–white-matter transition and underlying white matter, rather than representing a uniform increase in fiber-related signal throughout the cortical ribbon. We therefore interpret the cortical and subcortical profiles jointly: diffusivity-related differences extend broadly through the cortex, whereas fiber-related differences become increasingly expressed toward and beneath the gray–white matter interface.

### 2.3 The 3HG diffusion phenotype generalizes in vivo and varies with folding convergence

We first tested whether the tensor-derived diffusion pattern identified in fixed tissue was also present in the living brain. In 414 adults matched to the donor age range, FA was higher at 3HGs (*d* = 0.264), whereas MD (*d* = −0.220), RD (*d* = −0.252), and AD (*d* = −0.098) were lower (Fig. 3b). The same direction was observed in 90.8–99.3% of participants (all *P <* 10^*−*90^). When the *ex vivo* analysis was repeated using the same parcel-matched crown design, the diffusivity effects remained significant, whereas the FA difference was smaller (*d* = 0.058, *P* = 0.141). Thus, reduced cortical diffusivity showed particularly consistent agreement across the two datasets, while the FA effect was more sensitive to regional control selection.

**Fig. 3.**
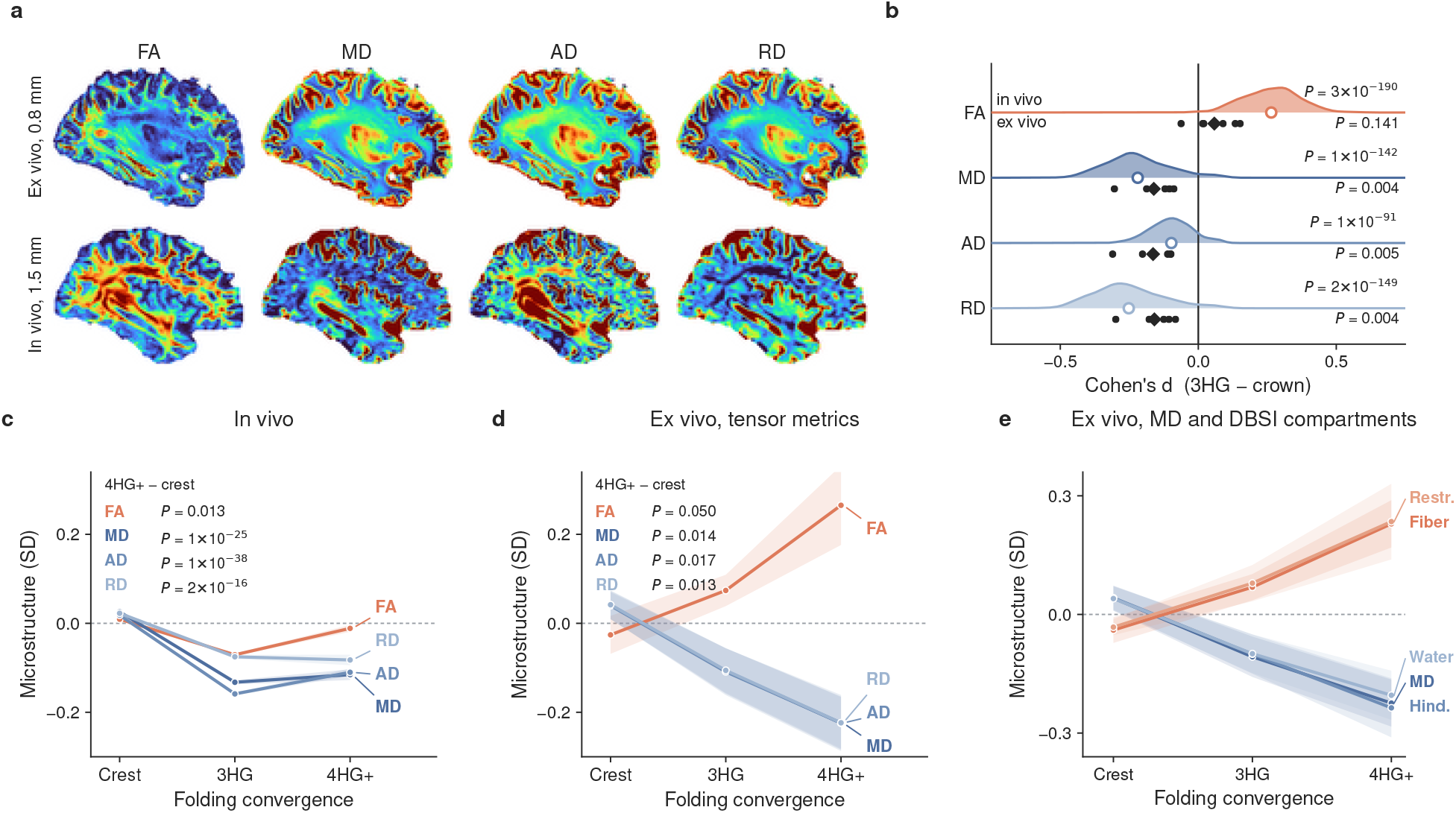
The cortical diffusion phenotype of three-hinge gyri is reproduced in vivo and varies with folding convergence. **a** Representative fractional anisotropy (FA), mean diffusivity (MD), axial diffusivity (AD), and radial diffusivity (RD) maps from the *ex vivo* 0.8-mm and *in vivo* 1.5-mm diffusion data. **b** 3HG–crown effect sizes for FA, MD, AD, and RD. For each metric, the shaded distribution above the horizontal reference line shows the subject-level *in vivo* effects across 414 adults, whereas the points below show the effects from the six *ex vivo* hemispheres. Positive Cohen’s *d* indicates higher values at 3HGs and negative values lower values. The same directional pattern is observed in both datasets, with higher FA and lower MD, AD, and RD at 3HGs, although the *ex vivo* FA difference is attenuated after parcel matching. For this cross-dataset comparison, crown controls in both datasets were selected using the same parcel-matched design. **c** *In vivo* diffusion measures across ordinary gyral crests, 3HGs, and higher-order junctions with degree *≥*4 (4HG+). The three folding classes were matched on their sulcal-depth distributions to separate folding convergence from position along the gyral–sulcal dimension. Values are standardized within participant. The major diffusivity shift occurs from ordinary crest to higher-order junctions, whereas the finer progression from 3HG to 4HG+ is less consistent *in vivo*. **d** Corresponding folding-convergence analysis for the tensor measures in the six *ex vivo* hemispheres. Increasing gyral convergence is accompanied by progressively lower MD, AD, and RD and higher FA. **e** Folding-convergence analysis for additional *ex vivo* diffusion measures. DBSI fiber and restricted fractions increase with gyral convergence, whereas MD, DBSI hindered fraction, and free-water fraction decrease. The crown controls in **b** and the skeleton-crest reference in **c–e** are distinct comparison groups: the former reproduces the primary 3HG–crown analysis, whereas the latter is restricted to the gyral skeleton and matched across folding classes on sulcal depth. FA, fractional anisotropy; MD, mean diffusivity; AD, axial diffusivity; RD, radial diffusivity; DBSI, diffusion basis spectrum imaging.

We then examined whether this phenotype extended beyond the categorical definition of a 3HG. In the *ex vivo* data, sulcal-depth-matched measurements showed a graded organization from ordinary gyral crest through 3HG to higher-order junctions (Fig. 3d,e). MD, AD, RD, DBSI hindered fraction, and free-water fraction decreased with increasing convergence, whereas FA, DBSI fiber fraction, and restricted fraction increased. The *in vivo* data reproduced the major diffusivity shift from ordinary gyral crest to higher-order junctions (Fig. 3c): MD, AD, and RD decreased markedly from crest to 3HG and remained lower at degree-≥ 4 junctions (4HG+). The finer progression beyond 3HG was not resolved in vivo, however, with MD and RD changing little and AD and FA showing non-monotonic profiles.

Within the *ex vivo* data, treating gyral-skeleton degree continuously yielded the same overall organization. After accounting for cortical region and sulcal depth, increasing degree was associated with lower MD, DBSI hindered fraction, and free-water fraction and with higher DBSI fiber fraction, restricted fraction, and FA. Sulcal-depth matching closely aligned the fold-position distributions across folding classes, and additional adjustment for local absolute curvature changed the degree slopes by less than 0.001 SD per order for every diffusion measure (Supplementary Fig. S4).

### 2.4 Three-hinge gyri show preferential inter-hinge structural wiring

Finally, we examined whether the folding and microstructural organization of 3HGs was accompanied by a characteristic structural wiring pattern. Structural connections were reconstructed using hemisphere-wide diffusion tractography, with nodes defined as superficial white-matter patches beneath each gyral junction and matched gyral-crown controls from the same cortical parcels (Fig. 4a,f). We first distinguished connection incidence from the streamline fraction carried by an existing connection. 3HG pairs showed a modestly higher incidence than crown pairs (1.14×, *P* = 0.035; Fig. 4b,c), whereas streamline fraction among connected pairs did not differ (0.96×, *P* = 0.34). This incidence difference was sensitive to gyral tractography bias. Endpoint density at the cortical positions occupied by 3HGs was 1.50× that at matched crown locations, and adjustment for the number of streamlines sampled at each node reduced the pair-incidence ratio to 1.02× (*P* = 0.45; Supplementary Fig. S2). Raw pair incidence therefore could not distinguish preferential wiring from differential streamline sampling. We therefore based the primary connectivity inference on a seed-normalized comparison in which 3HG and crown seeds were evaluated against the same 3HG target set. The fraction of sampled streamlines reaching another 3HG was 1.23× higher for 3HG seeds than for matched crown seeds (6/6 specimens, *P* = 0.010; Fig. 4d,e). Because the target set was identical for both seed classes and each seed was normalized by its own sampled streamline count, this contrast removes target-coverage differences and is not mechanically increased by a larger number of sampled streamlines. The corresponding seed-by-target interaction was 1.60× (*P* = 0.004), providing supporting evidence for preferential within-class wiring. Regional location and physical distance did not explain the observed pair-incidence difference. After matching 3HG and crown pairs by cortical-region pair and inter-node distance, the incidence ratio increased from 1.14× to 1.24× (6/6 specimens, *P* = 0.007; Fig. 4g). Because pair incidence remained sensitive to streamline sampling, however, this analysis was treated as a control rather than as the primary evidence for inter-hinge preference. The direction of the seed-normalized preference was preserved across the tested tractography parameter settings and alternative reconstruction strategies (Supplementary Figs. S1b and S3). Across parameter variations, the seed contrast remained above unity (1.06× –1.58×); changing the number of propagated fiber directions, the local direction-selection rule, or the tractography implementation likewise produced concordant enrichment. The magnitude of the effect therefore depended on tractography reconstruction, whereas its direction was preserved across the tested approaches.

**Fig. 4.**
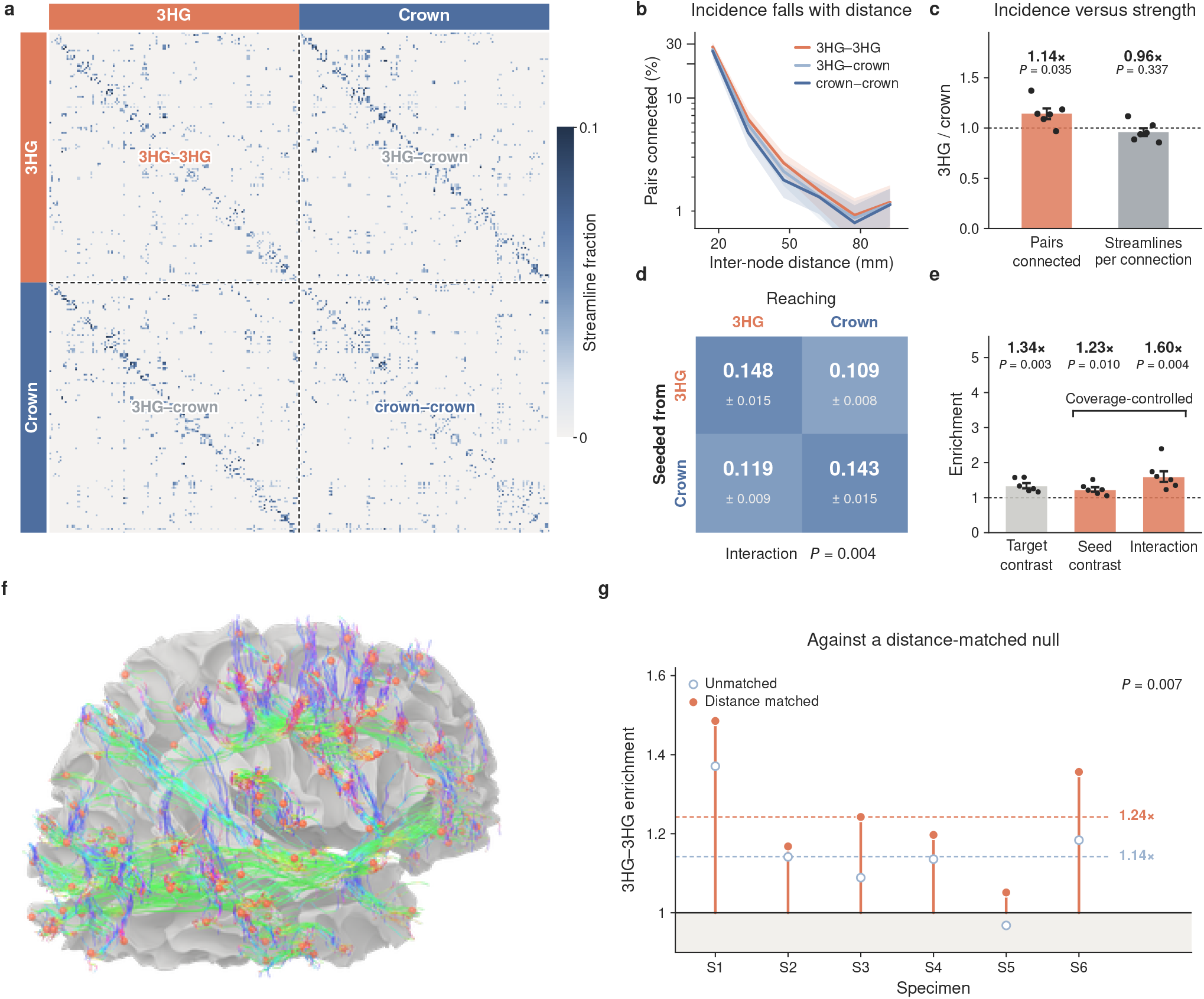
Three-hinge gyri show preferential inter-hinge structural wiring. **a** Representative node-to-node structural connectivity matrix from one *ex vivo* hemisphere. Rows denote seed nodes and columns denote target nodes. Each matrix element gives the fraction of streamlines sampled from a seed whose distal endpoint reached the corresponding target, with nodes grouped into 3HG and crown classes. **b** Probability that a node pair is connected as a function of inter-node distance for 3HG–3HG, 3HG–crown, and crown–crown pairs. Connection incidence decreases strongly with physical separation, motivating explicit control for distance in comparisons between node classes. **c** Raw 3HG–3HG relative to crown–crown enrichment separated into connection incidence and streamline fraction conditional on an existing connection. 3HG pairs show a modest increase in connection incidence (1.14×, P = 0.035), whereas the streamline fraction carried by connected pairs does not differ (0.96×, P = 0.337). **d** Mean seed-normalized streamline fractions for the four combinations of 3HG and crown seeds and targets. Each value represents the fraction of streamlines sampled from a seed class that reaches the indicated target class. **e** Enrichment derived from the four seed–target combinations in **d**. The target contrast compares 3HG and crown targets for 3HG seeds. The seed contrast compares 3HG and crown seeds against the same 3HG target set, thereby holding target coverage constant while normalizing each seed by its own sampled streamline count. Streamlines sampled beneath 3HGs are 1.23-fold more likely to reach another 3HG than streamlines sampled beneath matched crown controls (P = 0.010). The corresponding seed-by-target interaction is 1.60-fold (P = 0.004), providing additional evidence for preferential within-class wiring. **f** Representative inter-hinge tractography network from one right hemisphere. Orange spheres indicate 3HG nodes, and displayed streamlines connect distinct 3HGs and are colored according to local orientation. **g** 3HG–3HG connection-incidence enrichment relative to gyral-crown pairs before and after matching for cortical-region pair and inter-node distance. Distance and regional matching increase the across-specimen enrichment from 1.14× to 1.24× (P = 0.007).

## 3 Discussion

In this study, ultra-high-resolution *ex vivo* 7 T diffusion MRI revealed a consistent low-diffusivity, higher-anisotropy cortical profile at 3HGs. The diffusivity difference extended across the cortical ribbon, whereas anisotropy- and fiber-related differences became stronger toward the gray–white matter boundary and superficial white matter. The principal tensor-derived pattern was independently reproduced in 414 living adults. The *ex vivo* data further resolved variation with gyral-skeleton degree, while tractography showed preferential inter-hinge wiring among 3HGs.

### 3.1 Cortical diffusion microstructure and folding convergence

3HGs were characterized by lower MD, AD, and RD than ordinary gyral crowns, indicating reduced apparent water mobility within the cortical tissue. The accompanying reductions in DBSI hindered and free-water fractions further point to a distinct local diffusion environment, although these measures cannot be assigned to a specific cellular substrate. Cortical diffusion reflects contributions from cytoarchitecture, neurite packing and orientation, extracellular space, and myeloarchitecture [33–35]. The cortical-depth profiles refine this interpretation. MD, hindered fraction, and free-water fraction remained lower across much of the cortical ribbon, whereas FA and DBSI fiber fraction became more distinct toward the gray–white matter boundary. This difference is consistent with the changing balance of radial and tangential neurite orientations across cortical depth [33, 34, 36]. The 3HG phenotype therefore appears to contain a broadly intracortical low-diffusivity component and a more depth-dependent anisotropy-related component near the cortical–white-matter transition. The persistence of the diffusivity effects after restricting the analysis to high-gray-matter voxels further argues against simple white-matter partial volume as their primary source.

The cross-dataset comparison further distinguished these components. MD, AD, and RD remained lower at 3HGs under the same parcel-matched design in fixed and living tissue, whereas FA was less concordant in magnitude: the *ex vivo* effect became small after regional matching, while the *in vivo* contrast remained pronounced. One possible explanation follows directly from the cortical-depth profiles. The 3HG–crown FA difference was strongest toward the gray–white matter boundary rather than uniformly expressed through the cortex. At the 1.5-mm resolution of the *in vivo* data, cortical measurements integrate signal over a larger fraction of the cortical ribbon and its adjacent white matter, and may therefore incorporate more of this deeper anisotropic component. FA is also particularly sensitive to orientation dispersion and to the coexistence of radial and tangential neurite populations within a voxel [36]. Tissue state and acquisition provide additional sources of difference: fixation alters water mobility, relaxation, membrane permeability, and diffusion anisotropy [37–39], while DW-SSFP samples diffusion contrast differently from conventional *in vivo* encoding [40, 41]. These factors do not directly explain the smaller standardized FA effect *ex vivo*, but they make FA magnitude less comparable across datasets. In contrast, the consistent reduction of MD, AD, and RD identifies lower diffusivity as the more robust cross-dataset feature of the 3HG cortical phenotype.

The folding-convergence analysis places this phenotype beyond the categorical definition of a 3HG. In the *ex vivo* data, increasing gyral-skeleton degree was associated with progressively lower diffusivity and greater fiber-related diffusion after accounting for cortical region, sulcal depth, and local curvature. 3HGs therefore appear to occupy one position along a broader relationship between gyral convergence and local tissue organization, rather than forming an isolated microstructural class. Such a relationship is compatible with developmental accounts in which cortical expansion, cytoarchitectural differentiation, and the organization of underlying axonal pathways evolve in close interaction during gyrification [42–45]. The *in vivo* data reproduced the prominent diffusivity shift from ordinary gyral crest to higher-order junctions, but not the finer monotonic progression beyond 3HG observed *ex vivo*. FA showed an even less monotonic pattern across junction degrees *in vivo*. This is not necessarily inconsistent with the convergence effect: increasing junction complexity may simultaneously increase directional organization in some tissue components and orientation dispersion or crossing in others, influences that can act in opposite directions on a single-tensor FA measure. Combined with greater spatial averaging *in vivo*, this makes FA less suited than diffusivity to resolving subtle differences among higher-order junctions. The broad crest-to-junction organization therefore appears reproducible in the living cortex, whereas the finer ordering with gyral-skeleton degree is currently supported most clearly by the submillimeter *ex vivo* data.

### 3.2 Superficial white matter and inter-hinge wiring

The fiber-related differences became most apparent around and beneath the white surface. This localization is consistent with the geometry of superficial white matter, where short association fibers, longer-range pathways, and projection fibers bend and fan as they approach the cortex. Histological and diffusion studies have shown that these trajectories vary systematically across gyral crowns, walls, and fundi, and that superficial white-matter geometry strongly constrains how fibers reach the cortical sheet [46–48]. The increasing separation of the 3HG and crown DBSI fiber-fraction profiles beneath the white surface therefore indicates that the tissue below higher-order gyral junctions differs from that below ordinary gyral crowns. Importantly, this observation is derived directly from the diffusion signal and does not depend on streamline reconstruction. The tractography results provide a complementary view of the same anatomical compartment. Previous work has described 3HGs as densely connected or hub-like locations [19, 20, 22], but raw streamline counts are difficult to interpret because tractography preferentially reaches gyral crowns and is strongly influenced by superficial white-matter geometry [46, 47]. We observed this bias directly: the modest excess in raw 3HG–3HG connection incidence was largely attenuated after accounting for differences in streamline sampling. The more informative result came from the seed-normalized comparison. When 3HG and crown seeds were evaluated against the same 3HG target set, streamlines sampled beneath 3HGs were consistently more likely to reach another 3HG.

This preference was preserved when the number of initial tracking directions was increased, when the local direction-selection rule was changed, and when tractography was reconstructed independently in MRtrix3. The effect size varied across reconstructions, which is expected for diffusion tractography [49, 50], but its direction remained stable. The result is therefore best understood as a relative preference for inter-hinge wiring rather than as a direct estimate of anatomical fiber number. The convergence of the two analyses is more informative than either measurement alone. DBSI fiber fraction indicates a local difference in the superficial white matter beneath 3HGs, while tractography indicates that pathways sampled from the same locations preferentially reach other gyral hinges. These measurements have different sources of bias and probe different properties of the tissue, yet they point to the same anatomical region. This provides a broader interpretation of the high connectivity previously reported at 3HGs: higher-order gyral junctions are embedded in superficial white matter that is itself differently organized. 3HGs can therefore be viewed as locations where folding geometry, cortical diffusion microstructure, and superficial white-matter architecture converge. Their significance is not limited to being identifiable junctions on the cortical surface. The continuous relationship with skeleton degree further suggests that individualized folding topology contains information about local tissue organization that is not fully captured by cortical parcellation or conventional folding measures.

### 3.3 Limitations

The fine-scale analyses were based on six postmortem right hemispheres, reflecting the practical trade-off between sample size and the spatial resolution achievable with human *ex vivo* diffusion MRI [36]. The replication of lower MD, AD, and RD at 3HGs in 414 living adults supports the generalizability of the principal 3HG–crown diffusivity pattern, but does not independently validate the *ex vivo*-only findings. In particular, the cortical-depth, superficial white-matter, tractography, and finer folding-convergence effects require replication in additional high-resolution datasets. The restriction to right hemispheres also leaves possible hemispheric differences unresolved. The *ex vivo* and *in vivo* measurements are not quantitatively interchangeable. Fixation alters relaxation, water mobility, and diffusion anisotropy, while DW-SSFP combines diffusion weighting over multiple coherence pathways and cannot be assigned a single physical *b* value in the same way as conventional PGSE acquisitions [51–53]. Absolute diffusion values and cross-dataset effect magnitudes should therefore be interpreted cautiously. The reproducibility of the relative MD, AD, and RD differences is more informative than agreement in their absolute values, while the less concordant FA effects likely reflect the greater sensitivity of anisotropy to tissue state, acquisition, spatial scale, and regional context. Finally, diffusion MRI and DBSI provide indirect measures of tissue organization and do not identify the cellular substrate of the 3HG phenotype. Registered histology will be required to relate these signals to neurite architecture, myeloarchitecture, and extracellular structure [54]. The observed association between folding convergence and diffusion organization is also cross-sectional and does not establish developmental causality; longitudinal developmental imaging and biomechanical modeling will be needed to determine how these features become coupled.

## 4 Methods

### 4.1 Postmortem specimens and 7T MRI

We analyzed six postmortem right cerebral hemispheres from a publicly available multimodal 7T MRI dataset [55]. The six postmortem hemispheres were obtained from the National Health and Disease Human Brain Tissue Repository. Donors or their legally authorized next-of-kin provided written informed consent for clinical autopsy and brain donation for research under a protocol approved by the Institutional Review Board of the original study [55]. Neuropathological examination reported no significant abnormalities. Following autopsy, the hemispheres were fixed in phosphate-buffered 6% paraformaldehyde. Before imaging, the specimens were rinsed and soaked in saline, transferred to specimen-specific containers, immersed in Fomblin for susceptibility matching, and vacuum-treated to remove residual air bubbles. All images were acquired on a 7T MRI system (MAGNETOM 7T, Siemens Health-care, Erlangen, Germany) equipped with a 32-channel head coil. High-resolution structural images were acquired using a three-dimensional variable-flip-angle T2-weighted turbo spin-echo sequence at 0.5 × 0.5 × 0.6 mm^3^. High-angular-resolution diffusion MRI (HARDI) was acquired using a three-dimensional diffusion-weighted steady-state free-precession (DW-SSFP) sequence at 0.8-mm isotropic resolution, with 20 non-diffusion-weighted acquisitions and 60 diffusion directions at a nominal diffusion weighting of *b* = 6000 s/mm^2^.

Diffusion preprocessing distributed with the dataset included denoising, Gibbs-ringing removal, and correction for motion and image distortion using MRtrix3 [56]. DW-SSFP data were analyzed according to the nominal diffusion weighting reported with the dataset. Diffusion tensor fitting provided mean diffusivity (MD), axial diffusivity (AD), radial diffusivity (RD), and fractional anisotropy (FA). Diffusion basis spectrum imaging (DBSI) [57] provided anisotropic fiber and isotropic hindered, restricted, and free-water fractions.

### 4.2 Cortical surface reconstruction and processing

White and pial cortical surfaces were reconstructed from the high-resolution postmortem T2-weighted images and accompanying tissue segmentations. Because fixed-tissue T2-weighted MRI does not provide the intensity ordering assumed by conventional T1-weighted surface-reconstruction procedures, the structural images were locally normalized and transformed to a T1-like contrast while retaining image information at the gray–white matter boundary (Fig. 5a,b). Initial meshes were generated from the tissue segmentation, topology corrected at reduced mesh resolution, and subsequently fitted to the image-derived gray–white boundary. Sulcal spaces that had collapsed during fixation were recovered before pial-surface estimation [58, 59]. Full reconstruction details are provided in the Supplementary S2.1. Surface orientation and handedness were standardized across specimens, followed by curvature-based spherical registration to a common cortical reference [60, 61]. Desikan–Killiany labels [62] were transferred through the spherical correspondence. Cortical thickness, sulcal depth, and curvature were derived from the reconstructed surfaces, with the medial wall excluded from all subsequent analyses (Fig. 5c).

**Fig. 5.**
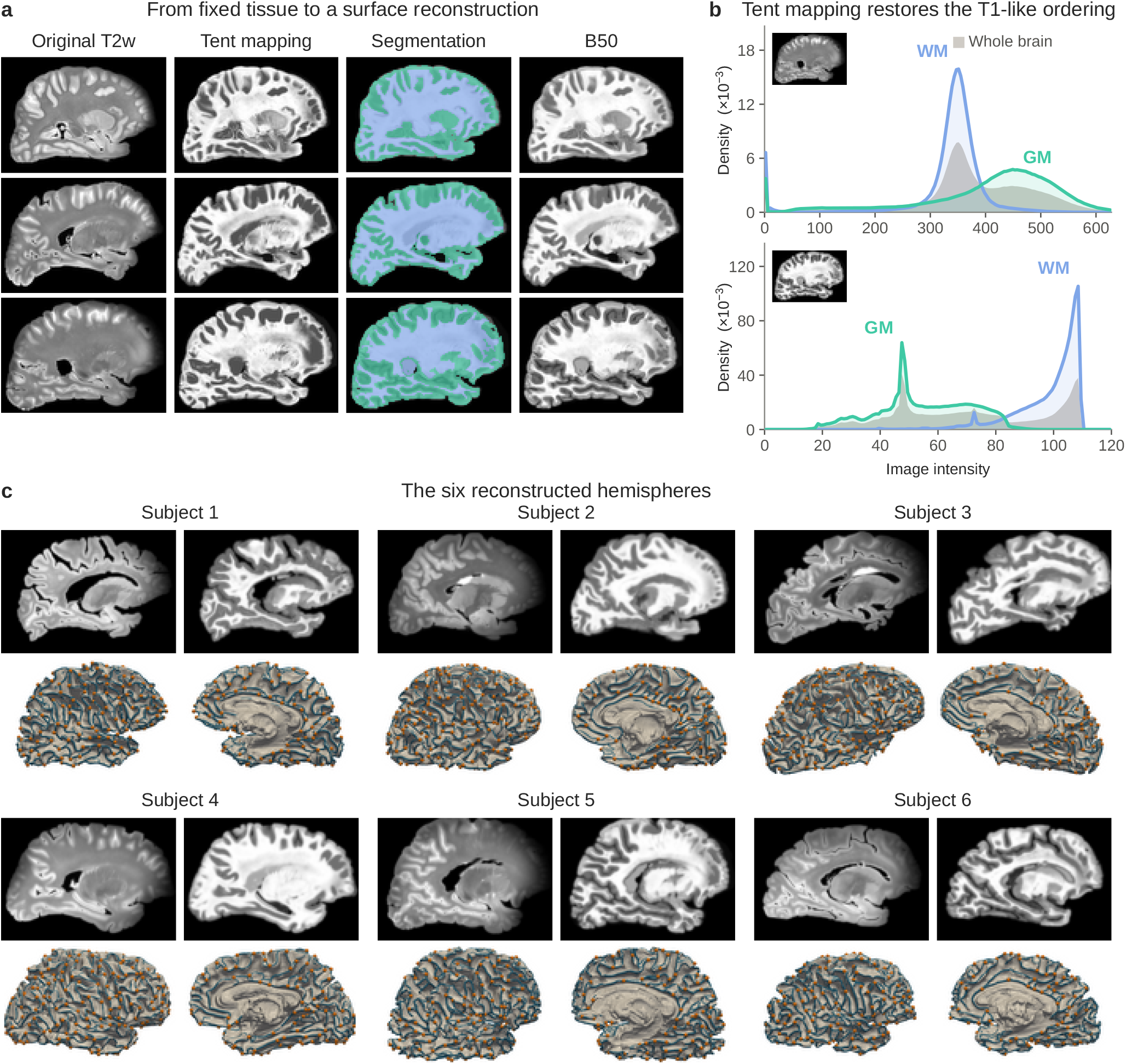
Postmortem cortical surface reconstruction and three-hinge gyral mapping. **a** Representative stages of structural processing in three specimens, including the original T2-weighted image, T1-like intensity mapping, tissue segmentation, and the combined B50 image used for surface placement. **b** Gray- and white-matter intensity distributions before and after T1-like contrast construction, showing restoration of the intensity ordering used for cortical surface reconstruction. **c** Structural images and reconstructed cortical surfaces for the six *ex vivo* right hemispheres. The surfaces were used for gyral-skeleton extraction, three-hinge gyral identification, sampling across cortical depth, and structural connectivity analysis.

3HGs were identified from the gyral skeleton using the automated procedure described previously [27]. Briefly, gyral regions were defined from sulcal depth and curvature, thinned to obtain central gyral ridges, and represented as a surface graph. Skeleton vertices with at least three incident branches were classified as higher-order gyral junctions. Degree-3 junctions were designated 3HGs, whereas junctions of degree ≥ 4 were treated as higher-order junctions in the folding-convergence analysis. Spatially adjacent detections belonging to the same anatomical junction were aggregated before inference. For the cortical diffusion analyses, detections within 4 mm were represented by a single junction. Ordinary gyral crowns served as the principal reference class and were restricted to the most gyral quartile of the sulcal-depth distribution. Crown samples overlapping the neighborhood of a 3HG were excluded. Sulcal fundi, defined from the most sulcal quartile, were used as an additional anatomical reference. Depending on the analysis, crown samples were either count matched to 3HGs or selected within the same DK region.

### 4.3 Cortical diffusion analysis

Volumetric MRI measures were sampled within the cortical ribbon. The primary cortical analysis used measurements averaged over normalized depths 0.3–0.7 to reduce sensitivity to the white and pial boundaries. For whole-hemisphere comparisons, equal numbers of crown locations were randomly drawn from the eligible crown pool, with 20 repeated draws used to reduce dependence on a particular control sample. The 3HG–crown difference for each metric was summarized within each specimen using Cohen’s *d*. To assess whether these effects reflected the nonuniform regional distribution of 3HGs, a complementary analysis compared 3HG and crown measurements within individual DK parcels. Class differences were standardized by the corresponding within-region variability, producing region-specific effect estimates.

Nine equivolumetric surfaces were generated between the white and pial boundaries at normalized cortical depths *ρ* = 0.1, …, 0.9 using the curvature-corrected formulation of [63]. MD, FA, and the four DBSI fractions were sampled at each depth. Statistical analysis of the depth profiles was performed within anatomical regions. For each specimen, metric, DK parcel, and cortical depth, the 3HG–crown difference was standardized by the corresponding within-parcel variability. Eligible parcel-level contrasts were averaged to obtain one region-adjusted depth profile per specimen. The depth-averaged contrast quantified the overall 3HG–crown difference across the cortical ribbon, whereas a class-by-depth interaction tested whether this difference varied systematically with cortical depth. To examine the transition into underlying white matter, DBSI fiber fraction was additionally sampled along the local surface normal from 2 mm above the white surface within cortex to 5 mm beneath it. Profiles were summarized separately for 3HG and crown locations and compared at the specimen level. Sensitivity to tissue partial volume was assessed independently as described in the Supplementary Methods.

### 4.4 Folding-convergence analysis

We examined whether the 3HG-associated phenotype formed part of a broader relationship between folding convergence and cortical microstructure. For visualization, gyral-skeleton locations were grouped as ordinary gyral crest, 3HGs, or 4HGs+. The three groups were matched on their sulcal-depth distributions before comparison. A complementary continuous analysis was performed over gyral-skeleton vertices with degree 2–6. Each microstructural measure was standardized within specimen and DK parcel, and the association with skeleton degree was estimated while adjusting for sulcal depth:

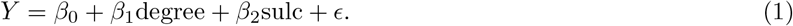

A second model additionally included local absolute curvature to test whether the degree effect could be explained by local surface bending. Models were fitted independently within each specimen, and specimen-specific degree coefficients were carried forward to group inference.

### 4.5 Structural connectivity analysis

Fiber orientation distributions were independently estimated from the preprocessed HARDI data using constrained spherical deconvolution. Deterministic hemisphere-wide tractography was then performed in DIPY [64], with white-matter seeding, a 0.4-mm tracking step, a 60^*°*^ maximum turning angle, and FA-based termination at 0.12. The resulting tractograms contained approximately 2.0–3.5 × 10^5^ streamlines per hemisphere. Detailed orientation-estimation and tracking parameters are provided in the Supplementary Methods. Connectivity was evaluated in superficial white matter beneath the cortical landmarks. Each retained 3HG node was paired with an eligible crown node from the same DK parcel. For each seed node *i*, connectivity to target *j* was expressed as the fraction of sampled streamlines whose distal endpoint reached the target neighborhood,

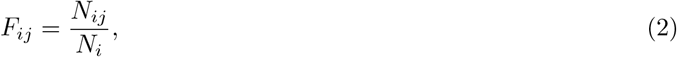

where *N*_*i*_ is the number of streamlines sampled by the seed and *N*_*ij*_ the number terminating near target *j*. Class-level connectivity was summarized for 3HG and crown seeds and targets. The primary measure was the seed contrast:

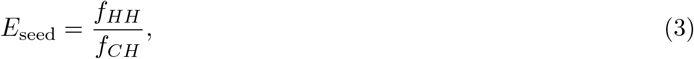

which compares 3HG and crown seeds against the same 3HG target set while normalizing each seed by its own sampled streamline count. Target and seed-by-target interaction contrasts were examined as complementary measures. Connection incidence and conditional connection strength were also quantified descriptively. Because tractography preferentially samples gyral locations [46, 47], gyral sampling bias was explicitly evaluated, and inferential interpretation focused on the seed-normalized contrast rather than raw pair incidence. Distance- and region-matched crown pairs provided an additional null model for the observed inter-hinge preference. Tractography parameter and reconstruction sensitivity analyses are described in the Supplementary Methods.

### 4.6 In vivo validation

An independent *in vivo* analysis was performed in 414 typically aging adults from HCPA/AABC Release 2 [65, 66] whose ages matched the *ex vivo* donor range (48–69 years). For each participant, the baseline imaging visit was used. Sessions flagged with any diffusion or structural QC issue code in the HCPA/AABC Release 2 release notes were excluded. All participants provided informed consent, and the HCP-A/AABC study was conducted with Institutional Review Board approval as described in the original study protocols [65, 66]. The final cohort comprised 244 women and 170 men (mean age, 58.1 years). Diffusion tensors were fitted using the nominal *b* = 0 volumes (*b* ≈ 5 s*/*mm^2^) and the complete *b* = 1500 s*/*mm^2^ shell (*b* = 1490–1510 s*/*mm^2^)of the preprocessed diffusion data using weighted least squares, yielding FA, MD, AD, and RD. The *b* = 3000 s*/*mm^2^ shell was not included in the tensor fit. Tensor maps were analyzed in the native T1w space to which the diffusion data had been registered. 3HGs and gyral-crown controls were defined using the same folding criteria as in the *ex vivo* analysis. Crown controls were selected from the most gyral quartile within the same DK parcel and were required to lie at least 8 mm geodesically from a 3HG. Twenty control draws were averaged for each subject. Because the *in vivo* diffusion data had lower spatial resolution, measurements were averaged within 3-mm geodesic neighborhoods at the cortical mid-thickness coordinate; cortical-depth profiling was not attempted. For the folding-convergence analysis, non-junction gyral crest, 3HG, and 4HG+ junctions were matched on their sulcal-depth distributions. Measurements were standardized within subject, and the analysis was repeated over 50 matched draws.

### 4.7 Statistics and Reproducibility

For the *ex vivo* analyses, the hemisphere was treated as the independent unit of analysis. For the *in vivo* validation, each participant contributed one subject-level estimate and was treated as the independent unit. In vivo 3HG– crown effect sizes were tested against zero across participants, and the proportion of participants showing the same direction of effect was also reported. Measurements from individual vertices, cortical locations, streamline pairs, and streamlines were first summarized within each specimen before group-level testing. Unless otherwise stated, two-sided tests were performed across the six specimen-level estimates. Connectivity ratios were log-transformed before testing. Benjamini–Hochberg correction was applied within each family of diffusion measures. Given the small sample size, we also report the consistency of effect direction across specimens, with exact sign-flip tests used as a complementary analysis where appropriate. For the *in vivo* folding analysis, the 4HG+–crest contrast was computed within each participant and tested against zero across participants.

## Data Availability

The *ex vivo* data used in this study are publicly available via Figshare at https://doi.org/10.6084/m9.figshare.28211843. Data from the HCPA/AABC project are publicly available at https://www.humanconnectome.org/study/hcp-lifespan-aging.

## Code Availability

The source code and implementation used in this study are publicly available at https://github.com/m1nhengChen/exvivo-cortical-folding.

## Author Contributions

**M.C**.: Conceptualization, Methodology, Validation, Formal analysis, Investigation, Data Curation, Writing - Original Draft, Visualization, Writing - Review & Editing; **X.D**.: Data Curation, Writing - Review & Editing; **J.C**.: Data Curation, Writing - Review & Editing; **D.H**.: Writing - Review & Editing; **T.L**.: Supervision, Writing - Review & Editing; **D.Z**.: Writing - Review & Editing, Supervision, Funding acquisition; **G.L**.: Resources, Data Curation, Writing - Review & Editing, Supervision, Funding acquisition.

## Acknowledgements

This work was supported by the National Institutes of Health (R01AG075582, R01EB037388 and R01NS128534).

## References

[1] Chen, H., Li, Y., Ge, F., Li, G., Shen, D., Liu, T.: Gyral net: A new representation of cortical folding organization. Medical image analysis 42, 14–25 (2017) 10.1016/j.media.2017.07.001

[2] Zhang, T., Chen, H., Razavi, M.J., Li, Y., Ge, F., Guo, L., Wang, X., Liu, T.: Exploring 3-hinge gyral folding patterns among hcp q3 868 human subjects. Human brain mapping 39(10), 4134–4149 (2018) 10.1002/hbm.24237

[3] Ronan, L., Fletcher, P.C.: From genes to folds: a review of cortical gyrification theory. Brain Structure and Function 220(5), 2475–2483 (2015) 10.1007/s00429-014-0961-z

[4] Fornito, A., Yücel, M., Wood, S., Stuart, G.W., Buchanan, J.-A., Proffitt, T., Anderson, V., Velakoulis, D., Pantelis, C.: Individual differences in anterior cingulate/paracingulate morphology are related to executive functions in healthy males. Cerebral cortex 14(4), 424–431 (2004) 10.1093/cercor/bhh004

[5] Willbrand, E.H., Ferrer, E., Bunge, S.A., Weiner, K.S.: Development of human lateral prefrontal sulcal morphology and its relation to reasoning performance. The Journal of Neuroscience 43(14), 2552–2567 (2023) 10.1523/JNEUROSCI.1745-22.2023

[6] Santacroce, F., Cachia, A., Fragueiro, A., Grande, E., Roell, M., Baldassarre, A., Sestieri, C., Committeri, G.: Human intraparietal sulcal morphology relates to individual differences in language and memory performance. Communications Biology 7(1), 520 (2024) 10.1038/s42003-024-06175-9

[7] Brun, L., Auzias, G., Viellard, M., Villeneuve, N., Girard, N., Poinso, F., Da Fonseca, D., Deruelle, C.: Localized misfolding within broca’s area as a distinctive feature of autistic disorder. Biological Psychiatry: Cognitive Neuroscience and Neuroimaging 1(2), 160–168 (2016) 10.1016/j.bpsc.2015.11.003

[8] MacKinley, M.L., Sabesan, P., Palaniyappan, L.: Deviant cortical sulcation related to schizophrenia and cognitive deficits in the second trimester. Translational neuroscience 11(1), 236–240 (2020) 10.1515/tnsci-2020-0111

[9] Shan, Y., Qiao, H., He, Y., Chu, L., Zeng, D., Dong, X., Zhao, T., Liao, X., Chen, X., Xia, Y., et al.: Linking changes in sulcal morphometry to cognitive development from childhood to adolescence. Communications Biology (2026) 10.1038/s42003-026-09956-6

[10] Cachia, A., Borst, G., Tissier, C., Fisher, C., Plaze, M., Gay, O., Riviére, D., Gogtay, N., Giedd, J., Mangin, J.-F., et al.: Longitudinal stability of the folding pattern of the anterior cingulate cortex during development. Developmental cognitive neuroscience 19, 122–127 (2016) 10.1016/j.dcn.2016.02.011

[11] Cachia, A., Borst, G., Jardri, R., Raznahan, A., Murray, G.K., Mangin, J.-F., Plaze, M.: Towards deciphering the fetal foundation of normal cognition and cognitive symptoms from sulcation of the cortex. Frontiers in Neuroanatomy 15, 712862 (2021) 10.3389/fnana.2021.712862

[12] Bok, S.T.: Der einflu der in den furchen und windungen auftretenden krümmungen der gro hirnrinde auf die rindenarchitektur. Zeitschrift für die gesamte Neurologie und Psychiatrie 121(1), 682–750 (1929) 10.1007/BF02864437

[13] Hilgetag, C.C., Barbas, H.: Developmental mechanics of the primate cerebral cortex. Anatomy and embryology 210(5), 411–417 (2005) 10.1007/s00429-005-0041-5

[14] Welker WPeters, A., Jones, E.: Why does cerebral cortex fissure and fold. A review of determinants of gyri and sulci Cerebral cortex, 3–136 (1990) 10.1007/978-1-4615-3824-0_1

[15] Nie, J., Guo, L., Li, K., Wang, Y., Chen, G., Li, L., Chen, H., Deng, F., Jiang, X., Zhang, T., et al.: Axonal fiber terminations concentrate on gyri. Cerebral cortex 22(12), 2831–2839 (2012) 10.1093/cercor/bhr361

[16] Chen, H., Zhang, T., Guo, L., Li, K., Yu, X., Li, L., Hu, X., Han, J., Hu, X., Liu, T.: Coevolution of gyral folding and structural connection patterns in primate brains. Cerebral Cortex 23(5), 1208–1217 (2013) 10.1093/cercor/bhs113

[17] Jiang, X., Li, X., Lv, J., Zhang, T., Zhang, S., Guo, L., Liu, T.: Sparse representation of hcp grayordinate data reveals novel functional architecture of cerebral cortex. Human brain mapping 36(12), 5301–5319 (2015) 10.1002/hbm.23013

[18] Li, K., Guo, L., Li, G., Nie, J., Faraco, C., Cui, G., Zhao, Q., Miller, L.S., Liu, T.: Gyral folding pattern analysis via surface profiling. NeuroImage 52(4), 1202–1214 (2010) 10.1016/j.neuroimage.2010.04.263

[19] Ge, F., Li, X., Razavi, M.J., Chen, H., Zhang, T., Zhang, S., Guo, L., Hu, X., Wang, X., Liu, T.: Denser growing fiber connections induce 3-hinge gyral folding. Cerebral cortex 28(3), 1064–1075 (2018) 10.1093/cercor/bhx227

[20] Zhang, T., Li, X., Jiang, X., Ge, F., Zhang, S., Zhao, L., Liu, H., Huang, Y., Wang, X., Yang, J., Guo, L., Hu, X., Liu, T.: Cortical 3-hinges could serve as hubs in cortico-cortical connective network. Brain imaging and behavior 14(6), 2512–2529 (2020) 10.1007/s11682-019-00204-6

[21] Li, X., Chen, H., Zhang, T., Yu, X., Jiang, X., Li, K., Li, L., Razavi, M.J., Wang, X., Hu, X., Han, J., Guo, L., Hu, X., Liu, T.: Commonly preserved and species-specific gyral folding patterns across primate brains. Brain structure and function 222(5), 2127–2141 (2017) 10.1007/s00429-016-1329-3

[22] He, Z., Du, L., Huang, Y., Jiang, X., Lv, J., Guo, L., Zhang, S., Zhang, T.: Gyral hinges account for the highest cost and the highest communication capacity in a corticocortical network. Cerebral Cortex 32(16), 3359–3376 (2022) 10.1093/cercor/bhab420

[23] Chen, M., Chen, T., Zhuang, Y., Cao, C., Zhang, J., Liu, T., Zhang, L., Zhu, D.: Community-level modeling of gyral folding patterns for robust and anatomically informed individualized brain mapping. NeuroImage, 122037 (2026) 10.1016/j.neuroimage.2026.122037

[24] Zhuang, Y., Chen, M., Cao, C., Chen, T., Zhang, J., Yu, X., Lyu, Y., Zhang, L., Liu, T., Zhu, D.: Gyralnet subnetwork partitioning via differentiable spectral modularity optimization. In: International Workshop on Machine Learning in Medical Imaging, pp. 477–486 (2025). 10.1007/978-3-032-09513-8_46. Springer

[25] Chen, T., Chen, M., Zhuang, Y., Zhang, J., Zhang, L., Liu, T., Blamire, A.M., O’Brien, J.T., Su, L., Zhu, D.: Representing the cognitive impairment continuum of alzheimer’s disease and lewy body dementia with a novel finer-scale cortical representation via disease embedding tree. Alzheimer’s & Dementia 21, 102179 (2025) 10.1002/alz70856_102179

[26] Chen, T., Chen, M., Zhang, J., Zhuang, Y., Cao, C., Yu, X., Lyu, Y., Zhang, L., Su, L., Liu, T., Zhu, D.: A unified continuous staging framework for alzheimer’s disease and lewy body dementia via hierarchical anatomical features. In: International Conference on Medical Image Computing and Computer-Assisted Intervention, pp. 13–23 (2025). 10.1007/978-3-032-04947-6_2. Springer

[27] Chen, M., Cao, C., Chen, T., Alhamad, D., Liu, T., Su, L., Zhu, D.: Fine-scale individualized gyral folding-based cortical similarity networks reveal distinct organizational patterns in alzheimer’s disease and lewy body dementia. Imaging Neuroscience 4, 1322 (2026) 10.1162/IMAG.a.1322

[28] Casamitjana, A., Mancini, M., Robinson, E., Peter, L., Annunziata, R., Althonayan, J., Crampsie, S., Blackburn, E., Billot, B., Atzeni, A., et al.: A probabilistic histological atlas of the human brain for mri segmentation. Nature 648(8094), 678–685 (2025) 10.1038/s41586-025-09708-2

[29] Yushkevich, P.A., Muñoz López, M., Onzoño Martin, M.M., Ittyerah, R., Lim, S., Ravikumar, S., Bedard, M.L., Pickup, S., Liu, W., Wang, J., et al.: Three-dimensional mapping of neurofibrillary tangle burden in the human medial temporal lobe. Brain 144(9), 2784–2797 (2021) 10.1093/brain/awab262

[30] Alkemade, A., Bazin, P.-L., Balesar, R., Pine, K., Kirilina, E., Möller, H.E., Trampel, R., Kros, J.M., Keuken, M.C., Bleys, R.L., et al.: A unified 3d map of microscopic architecture and mri of the human brain. Science advances 8(17), 7892 (2022) 10.1126/sciadv.abj7892

[31] Beaujoin, J., Palomero-Gallagher, N., Boumezbeur, F., Axer, M., Bernard, J., Poupon, F., Schmitz, D., Mangin, J.-F., Poupon, C.: Post-mortem inference of the human hippocampal connectivity and microstructure using ultra-high field diffusion mri at 11.7 t. Brain Structure and Function 223(5), 2157–2179 (2018) 10.1007/s00429-018-1617-1

[32] Ravikumar, S., Wisse, L.E., Lim, S., Ittyerah, R., Xie, L., Bedard, M.L., Das, S.R., Lee, E.B., Tisdall, M.D., Prabhakaran, K., et al.: Ex vivo mri atlas of the human medial temporal lobe: characterizing neurodegeneration due to tau pathology. Acta neuropathologica communications 9(1), 173 (2021) 10.1186/s40478-021-01275-7

[33] Leuze, C.W., Anwander, A., Bazin, P.-L., Dhital, B., Stüber, C., Reimann, K., Geyer, S., Turner, R.: Layer-specific intracortical connectivity revealed with diffusion mri. Cerebral cortex 24(2), 328–339 (2014) 10.1093/cercor/bhs311

[34] Aggarwal, M., Nauen, D.W., Troncoso, J.C., Mori, S.: Probing region-specific microstructure of human cortical areas using high angular and spatial resolution diffusion mri. Neuroimage 105, 198–207 (2015) 10.1016/j.neuroimage.2014.10.053

[35] Sadikov, A., Choi, H.L., Xiao, J., Cai, L.T., Mukherjee, P.: Mapping the microstructure of human cerebral cortex in vivo with diffusion mri. Communications Biology 8(1), 1088 (2025) 10.1038/s42003-025-08523-9

[36] Roebroeck, A., Miller, K.L., Aggarwal, M.: Ex vivo diffusion mri of the human brain: Technical challenges and recent advances. NMR in Biomedicine 32(4), 3941 (2019) 10.1002/nbm.3941

[37] Shepherd, T.M., Thelwall, P.E., Stanisz, G.J., Blackband, S.J.: Aldehyde fixative solutions alter the water relaxation and diffusion properties of nervous tissue. Magnetic Resonance in Medicine: An Official Journal of the International Society for Magnetic Resonance in Medicine 62(1), 26–34 (2009) 10.1002/mrm.21977

[38] Shepherd, T.M., Flint, J.J., Thelwall, P.E., Stanisz, G.J., Mareci, T.H., Yachnis, A.T., Blackband, S.J.: Postmortem interval alters the water relaxation and diffusion properties of rat nervous tissue—implications for mri studies of human autopsy samples. Neuroimage 44(3), 820–826 (2009) 10.1016/j.neuroimage.2008.09.054

[39] Richardson, S., Siow, B., Panagiotaki, E., Schneider, T., Lythgoe, M.F., Alexander, D.C.: Viable and fixed white matter: diffusion magnetic resonance comparisons and contrasts at physiological temperature. Magnetic resonance in medicine 72(4), 1151–1161 (2014) 10.1002/mrm.25012

[40] McNab, J.A., Miller, K.L.: Sensitivity of diffusion weighted steady state free precession to anisotropic diffusion. Magnetic Resonance in Medicine: An Official Journal of the International Society for Magnetic Resonance in Medicine 60(2), 405–413 (2008) 10.1002/mrm.21668

[41] McNab, J.A., Miller, K.L.: Steady-state diffusion-weighted imaging: theory, acquisition and analysis. NMR in Biomedicine 23(7), 781–793 (2010) 10.1002/nbm.1509

[42] Garcia, K., Kroenke, C., Bayly, P.: Mechanics of cortical folding: stress, growth and stability. Philosophical Transactions of the Royal Society B: Biological Sciences 373(1759), 20170321 (2018) 10.1098/rstb.2017.0321

[43] Fernández, V., Borrell, V.: Developmental mechanisms of gyrification. Current Opinion in Neurobiology 80, 102711 (2023) 10.1016/j.conb.2023.102711

[44] Akula, S.K., Exposito-Alonso, D., Walsh, C.A.: Shaping the brain: The emergence of cortical structure and folding. Developmental cell 58(24), 2836–2849 (2023) 10.1016/j.devcel.2023.11.004

[45] Normand, F., Gajwani, M., Cao, T., Cruddas, J., Sangchooli, A., Oldham, S., Holmes, A., Robinson, P.A., Pang, J.C., Fornito, A.: Geometric constraints on the architecture of mammalian cortical connectomes. Cell (2026) 10.1016/j.cell.2026.05.048

[46] Reveley, C., Seth, A.K., Pierpaoli, C., Silva, A.C., Yu, D., Saunders, R.C., Leopold, D.A., Ye, F.Q.: Superficial white matter fiber systems impede detection of long-range cortical connections in diffusion mr tractography. Proceedings of the National Academy of Sciences 112(21), 2820–2828 (2015) 10.1073/pnas.1418198112

[47] Schilling, K., Gao, Y., Janve, V., Stepniewska, I., Landman, B.A., Anderson, A.W.: Confirmation of a gyral bias in diffusion mri fiber tractography. Human Brain Mapping 39(3), 1449–1466 (2018) 10.1002/hbm.23936

[48] Cottaar, M., Bastiani, M., Boddu, N., Glasser, M.F., Haber, S., Van Essen, D.C., Sotiropoulos, S.N., Jbabdi, S.: Modelling white matter in gyral blades as a continuous vector field. NeuroImage 227, 117693 (2021) 10.1016/j.neuroimage.2020.117693

[49] Smith, R.E., Tournier, J.-D., Calamante, F., Connelly, A.: Anatomically-constrained tractography: improved diffusion mri streamlines tractography through effective use of anatomical information. Neuroimage 62(3), 1924–1938 (2012) 10.1016/j.neuroimage.2012.06.005

[50] Maier-Hein, K.H., Neher, P.F., Houde, J.-C., Câté, M.-A., Garyfallidis, E., Zhong, J., Chamberland, M., Yeh, F.-C., Lin, Y.-C., Ji, Q., et al.: The challenge of mapping the human connectome based on diffusion tractography. Nature communications 8(1), 1349 (2017) 10.1038/s41467-017-01285-x

[51] Schilling, K.G., Grussu, F., Ianus, A., Hansen, B., Howard, A.F., Barrett, R.L., Aggarwal, M., Michielse, S., Nasrallah, F., Syeda, W., et al.: Considerations and recommendations from the ismrm diffusion study group for preclinical diffusion mri: Part 2—ex vivo imaging: Added value and acquisition. Magnetic resonance in medicine 93(6), 2535–2560 (2025) 10.1002/mrm.30435

[52] Schilling, K.G., Howard, A.F.D., Grussu, F., Ianus, A., Hansen, B., Barrett, R.L.C., Aggarwal, M., Michielse, S., Nasrallah, F., Syeda, W., Wang, N., Veraart, J., Roebroeck, A., Bagdasarian, A.F., Eichner, C., Sepehrband, F., Zimmermann, J., Soustelle, L., Bowman, C., Tendler, B.C., Hertanu, A., Jeurissen, B., Verhoye, M., Frydman, L., Looij, Y., Hike, D., Dunn, J.F., Miller, K., Landman, B.A., Shemesh, N., Anderson, A., McKinnon, E., Farquharson, S., Dell’Acqua, F., Pierpaoli, C., Drobnjak, I., Leemans, A., Harkins, K.D., Descoteaux, M., Xu, D., Huang, H., Santin, M.D., Grant, S.C., Obenaus, A., Kim, G.S., Wu, D., Le Bihan, D., Blackband, S.J., Ciobanu, L., Fieremans, E., Bai, R., Leergaard, T.B., Zhang, J., Dyrby, T.B., Johnson, G.A., Cohen-Adad, J., Budde, M.D., Jelescu, I.O.: Considerations and recommendations from the ismrm diffusion study group for preclinical diffusion mri: Part 3—ex vivo imaging: Data processing, comparisons with microscopy, and tractography. Magnetic Resonance in Medicine 93(6), 2561–2582 (2025) 10.1002/mrm.30424

[53] Tendler, B.C., Foxley, S., Cottaar, M., Jbabdi, S., Miller, K.L.: Modeling an equivalent b-value in diffusion-weighted steady-state free precession. Magnetic Resonance in Medicine 84(2), 873–884 (2020) 10.1002/mrm.28169

[54] Zhao, Z., Cao, Z., Zhu, Q., Xu, H., Li, S., Zhu, L., Xu, G., Zhu, K., Zhang, J., Wu, D.: Layer-dependent effect of aβ-pathology on cortical microstructure with ex vivo human brain diffusion mri at 7 tesla. Human Brain Mapping 46(7) (2025) 10.1002/hbm.70222

[55] Zhu, Q., Li, S., Cao, Z., Shen, Y., Xu, H., Xu, G., Li, H., Cui, Z., Zhu, K., Zhao, Z., Zhang, J., Wu, D.: 7 tesla multimodal mri dataset of ex-vivo human brain. Scientific Data 12(1), 845 (2025) 10.1038/s41597-025-04932-x

[56] Tournier, J.-D., Smith, R., Raffelt, D., Tabbara, R., Dhollander, T., Pietsch, M., Christiaens, D., Jeurissen, B., Yeh, C.-H., Connelly, A.: Mrtrix3: A fast, flexible and open software framework for medical image processing and visualisation. Neuroimage 202, 116137 (2019) 10.1016/j.neuroimage.2019.116137

[57] Sun, P., George, A., Perantie, D.C., Trinkaus, K., Ye, Z., Naismith, R.T., Song, S.-K., Cross, A.H.: Diffusion basis spectrum imaging provides insights into ms pathology. Neurology: Neuroimmunology & Neuroinflammation 7(2), 655 (2019) 10.1212/NXI.0000000000000655

[58] Wang, L., Wu, Z., Chen, L., Sun, Y., Lin, W., Li, G.: ibeat v2. 0: a multisite-applicable, deep learning-based pipeline for infant cerebral cortical surface reconstruction. Nature protocols 18(5), 1488–1509 (2023) 10.1038/s41596-023-00806-x

[59] Dale, A.M., Fischl, B., Sereno, M.I.: Cortical surface-based analysis: I. segmentation and surface reconstruction. Neuroimage 9(2), 179–194 (1999) 10.1006/nimg.1998.0395

[60] Fischl, B., Sereno, M.I., Tootell, R.B., Dale, A.M.: High-resolution intersubject averaging and a coordinate system for the cortical surface. Human brain mapping 8(4), 272–284 (1999) https://doi.org/(SICI)1097-0193(1999)8:4?272::AID-HBM10?3.0.CO;2-4

[61] Fischl, B., Sereno, M.I., Dale, A.M.: Cortical surface-based analysis: Ii: inflation, flattening, and a surface-based coordinate system. Neuroimage 9(2), 195–207 (1999) 10.1006/nimg.1998.0396

[62] Desikan, R.S., Ségonne, F., Fischl, B., Quinn, B.T., Dickerson, B.C., Blacker, D., Buckner, R.L., Dale, A.M., Maguire, R.P., Hyman, B.T., et al.: An automated labeling system for subdividing the human cerebral cortex on mri scans into gyral based regions of interest. Neuroimage 31(3), 968–980 (2006) 10.1016/j.neuroimage.2006.01.021

[63] Waehnert, M.D., Dinse, J., Weiss, M., Streicher, M.N., Waehnert, P., Geyer, S., Turner, R., Bazin, P.-L.: Anatomically motivated modeling of cortical laminae. NeuroImage 93, 210–220 (2014) 10.1016/j.neuroimage.2013.03.078. In-vivo Brodmann Mapping of the Human Brain

[64] Garyfallidis, E., Brett, M., Amirbekian, B., Rokem, A., Van Der Walt, S., Descoteaux, M., Nimmo-Smith, I.: Dipy, a library for the analysis of diffusion mri data. Frontiers in Neuroinformatics Volume 8 - 2014 (2014) 10.3389/fninf.2014.00008

[65] Bookheimer, S.Y., Salat, D.H., Terpstra, M., Ances, B.M., Barch, D.M., Buckner, R.L., Burgess, G.C., Curtiss, S.W., Diaz-Santos, M., Elam, J.S., et al.: The lifespan human connectome project in aging: an overview. Neuroimage 185, 335–348 (2019) 10.1016/j.neuroimage.2018.10.009

[66] Harms, M.P., Somerville, L.H., Ances, B.M., Andersson, J., Barch, D.M., Bastiani, M., Bookheimer, S.Y., Brown, T.B., Buckner, R.L., Burgess, G.C., et al.: Extending the human connectome project across ages: Imaging protocols for the lifespan development and aging projects. Neuroimage 183, 972–984 (2018) 10.1016/j.neuroimage.2018.09.060

