## Supplementary information for "Three-hinge gyri link individualized folding topology to cortical and superficial white-matter diffusion organization"

### *Three-hinge gyri show reproducible cortical diffusion organization across depth and folding convergence*


October 6, 2026

#### S1 Supplementary Figures

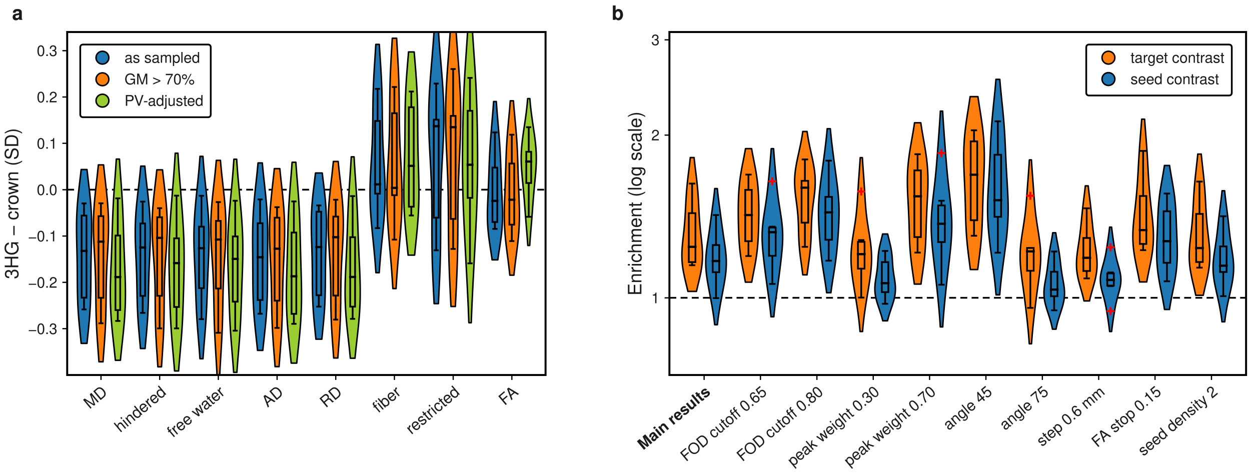


Figure S1: **Robustness of the intracortical diffusion and inter-hinge connectivity findings. a** Standardized 3HG–crown differences averaged across the nine equivolumetric cortical depths under three sampling strategies: the primary analysis, restriction to diffusion voxels containing at least 70% cortical gray matter, and explicit adjustment for white-matter and non-brain partial-volume fractions. The principal MD, DBSI hindered, and free-water effects are preserved or slightly increased after partial-volume control. **b** Target- and seed-based inter-hinge enrichment across ten tractography parameter settings. Each alternative differs from the reference reconstruction in one tracking parameter. Small points denote individual specimens and large points the geometric mean. Enrichment remains above unity across all tested settings, with the weakest seed contrast under the most permissive turning-angle criterion.

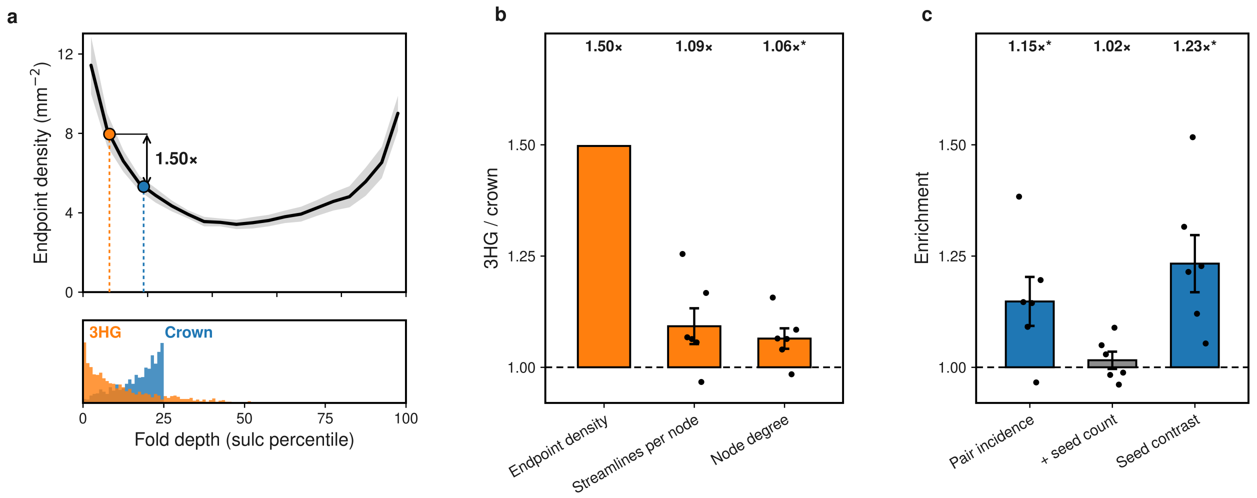


Figure S2: **Propagation of gyral tractography bias through the connectivity analysis.** **a** Area-normalized streamline endpoint density across cortical fold depth. The median folding positions occupied by 3HG and matched crown nodes differ by approximately 1.50-fold in endpoint density. **b** Attenuation of this difference from cortical endpoint density to the number of streamlines sampled by each sub-white-matter node and subsequently to graph node degree. **c** Sensitivity of inter-hinge enrichment to the connectivity estimator. The modest raw pair-incidence excess is strongly attenuated after adjustment for seed streamline count, whereas the primary seed contrast remains above unity. The latter compares both seed classes against the same 3HG target set and normalizes each seed by its own sampled streamline count.

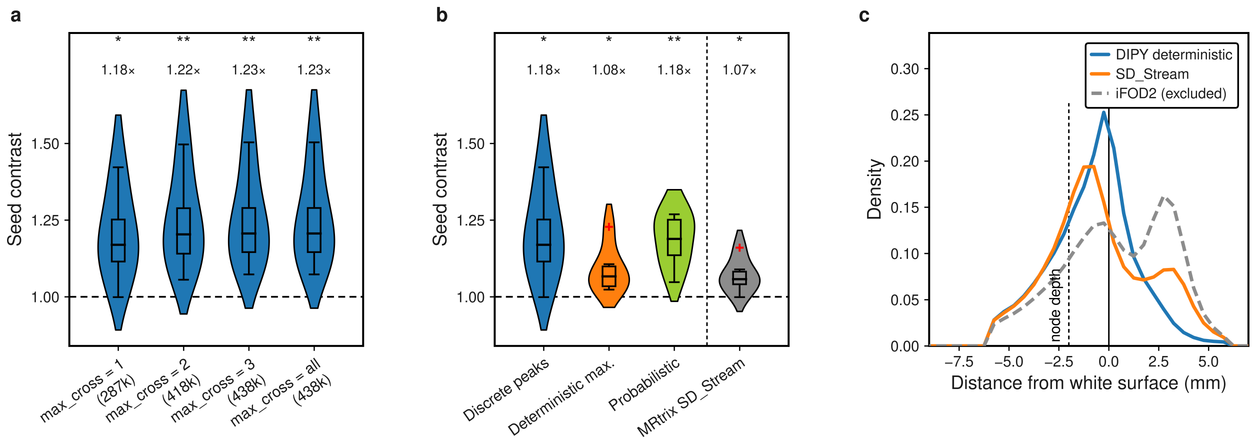


Figure S3: **Sensitivity of inter-hinge connectivity to tractography reconstruction.** **a** Seed-normalized inter-hinge contrast as the maximum number of initial tracking directions per seed (`max_cross`) was increased from one to all admissible FOD peaks. Violins show the distribution across six specimens; inner boxes indicate the median and interquartile range, whiskers indicate 1.5 × IQR, and red crosses denote values beyond the whiskers. **b** Seed contrast across alternative streamline reconstruction strategies. The first three reconstructions used the same CSD FOD, white-matter seeds, FA stopping criterion, maximum turning angle, and step size, differing only in the local direction-selection rule: discrete-peak tracking, deterministic propagation along the local FOD maximum, and probabilistic FOD sampling. MRtrix3 `SD_Stream` provided an independent deterministic reconstruction using a separately estimated response function and FOD. **c** Distribution of streamline endpoints as a function of signed distance from the white surface, with positive values directed toward the cortical ribbon. The dashed line marks the depth of the sub-WM node centers. MRtrix3 iFOD2 is shown for comparison but was excluded from the connectivity contrast because its endpoint distribution extended substantially beyond the gray–white matter interface, making it incompatible with the endpoint-based target definition used here. Geometric mean ratios are shown above the violins; $*P* < 0.05, **P* < 0.01$, two-sided one-sample t-tests on log ratios.

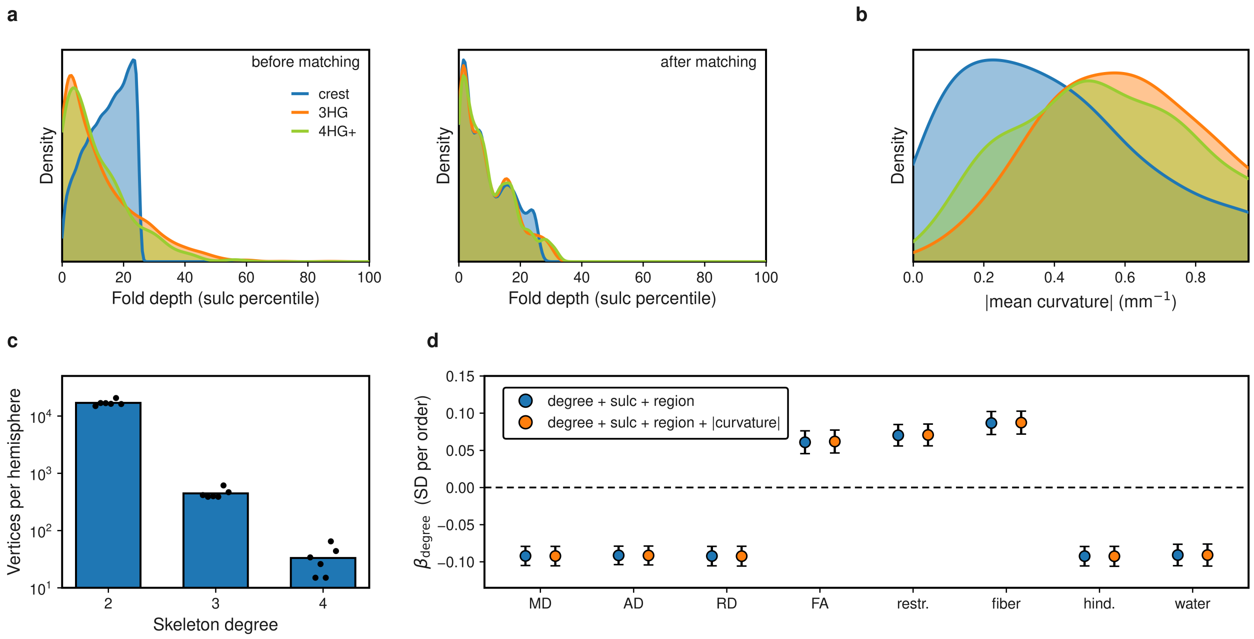


Figure S4: **Folding-convergence effects are preserved after matching for fold depth and adjustment for local curvature. a** Sulcal-depth percentile distributions for ordinary gyral crest, 3HGs, and higher-order junctions before and after matching. Vertices are pooled across the six hemispheres for visualization only. After matching, the three classes showed nearly identical fold-depth distributions (median percentile, 7.1, 7.1, and 7.3, respectively). **b** Distribution of absolute mean curvature after sulcal-depth matching. Curvature was not included in the matching criterion because it is intrinsically related to local junction geometry; its contribution was instead evaluated explicitly in the continuous degree model shown in **d**. **c** Number of gyral-skeleton vertices at each skeleton degree in the six hemispheres. Bars show the mean across hemispheres and points individual specimens; the ordinate is logarithmic. **d** Association between gyral-skeleton degree and each diffusion measure. Blue points show the mean hemisphere-level degree coefficient from models adjusting for sulcal depth and DK parcel; orange points show the corresponding coefficients after additional adjustment for absolute mean curvature. Error bars denote s.e.m. across six hemispheres. All eight diffusion measures retained a significant degree association ($P = 0.0007$–$0.011$), and curvature adjustment changed no mean slope by more than 0.001 SD per degree.

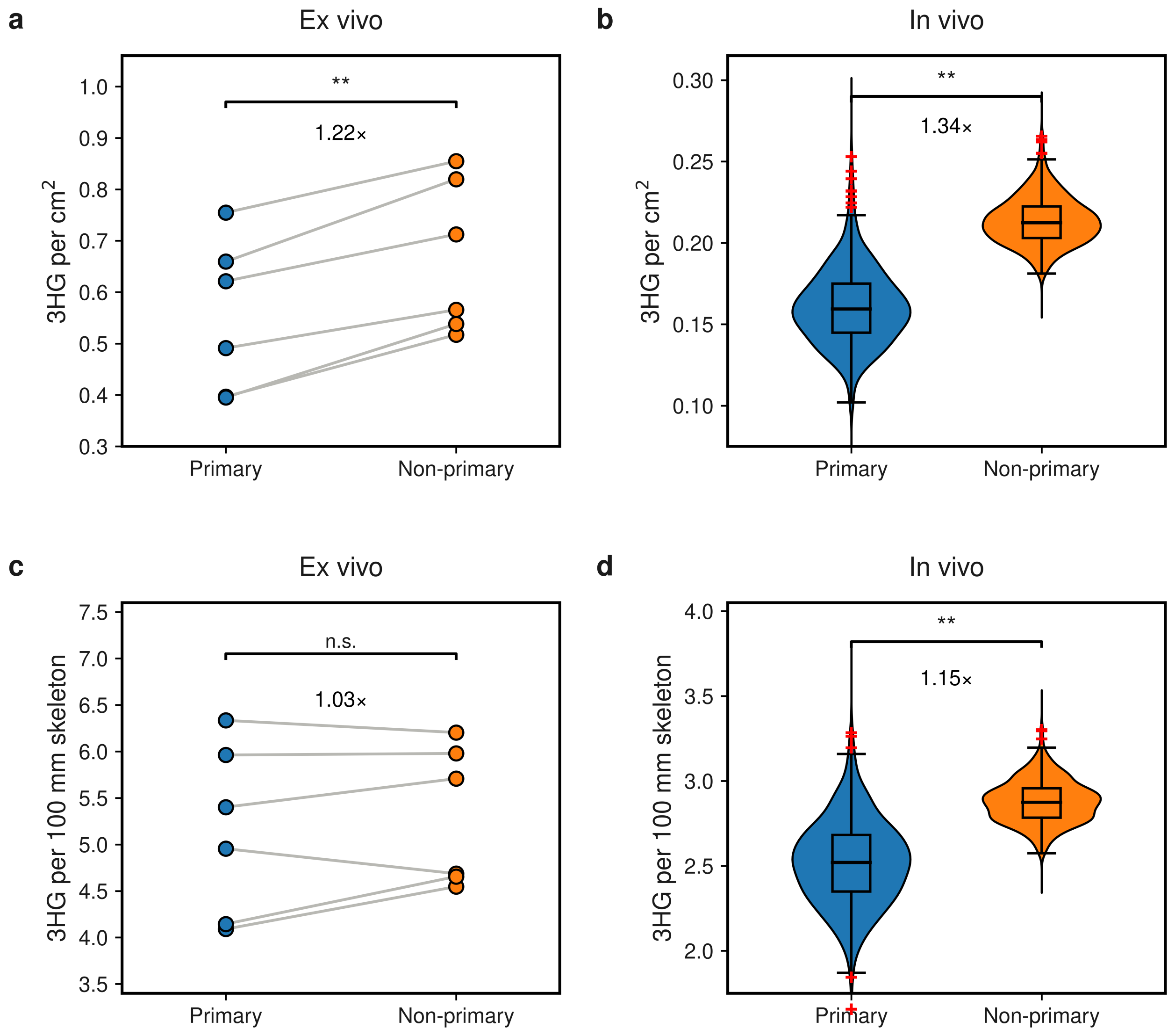


Figure S5: **Regional distribution of three-hinge gyri in primary and non-primary cortex.** **a** Surface-area-normalized 3HG density in the six *ex vivo* right hemispheres. 3HG density was 1.22-fold higher in non-primary cortex (6/6 hemispheres; two-sided one-sample $t$-test of the log density ratio, $t(5) = 6.35$, $P = 0.0014$). **b** Corresponding analysis in 414 *in vivo* participants, with both hemispheres pooled within participant. Non-primary cortex showed 1.34-fold higher 3HG density (95% CI, 1.32–1.36; $t(413) = 42.6$, $P = 2 \times 10^{-153}$), with the same direction observed in 98.8% of participants. **c** *Ex vivo* sensitivity analysis after normalizing the number of 3HGs by total gyral-skeleton length rather than cortical surface area. The non-primary-to-primary ratio was reduced to 1.03 (95% CI, 0.96–1.12; $t(5) = 1.19$, $P = 0.29$). **d** Corresponding gyral-skeleton-length-normalized analysis *in vivo*. The non-primary enrichment remained present (1.15-fold; 95% CI, 1.14–1.16; $t(413) = 27.9$, $P = 7 \times 10^{-97}$), with 91.5% of participants showing the same direction. Surface-area and skeleton-length normalization used the same analyzable cortical regions and 3HG detection criteria within each dataset. $^{**}P < 0.01$; n.s., not significant.

### S2 Supplementary Methods

#### S2.1 Detailed cortical surface reconstruction

Cortical surfaces were reconstructed directly from the high-resolution postmortem MRI rather than transferred from an external template. The reconstruction pipeline was adapted to two properties of the dataset that differ substantially from conventional *in vivo* T1-weighted imaging: the presence of an open medial cutting plane in the isolated hemispheres and the altered GM–WM contrast of fixed-tissue T2-weighted MRI. The procedure consisted of volumetric mask preparation, construction of an image with T1-like GM–WM intensity ordering, white-surface reconstruction, and recovery of collapsed sulcal spaces before pial-surface placement.

##### S2.1.1 Volume preparation

Cortical gray-matter (GM) and white-matter (WM) segmentations distributed with the dataset were used to initialize the reconstruction. Because each specimen consisted of an isolated cerebral hemisphere, the medial cutting plane was open to the image exterior. Standard three-dimensional hole-filling operations are therefore not appropriate: internal cavities that are connected to the exterior through the medial cut can be incorrectly classified as background and remain unfilled, whereas unrestricted filling can introduce artificial connections across the cut plane. To distinguish enclosed WM cavities from regions connected to the image exterior, candidate voxels were evaluated along the six cardinal directions ($\pm x$, $\pm y$, and $\pm z$). For each candidate location, rays were followed toward the image boundary, and the presence of WM along each direction was recorded. Voxels surrounded by WM according to this directional criterion were treated as interior candidates, allowing local WM cavities to be repaired without closing the medial surface of the hemisphere. All subsequent surface reconstruction was performed on a common 0.5-mm isotropic working grid. To avoid staircase artifacts and label mixing introduced by direct interpolation of categorical segmentations, the GM and WM masks were first converted to signed-distance representations. The distance fields were resampled to the target grid and converted back to binary tissue masks using their zero level sets. This preserved the location of tissue boundaries more accurately than interpolation of the discrete labels themselves.

##### S2.1.2 Image-derived T1-like contrast

Fixed-tissue T2-weighted MRI does not exhibit the WM–GM intensity ordering assumed by conventional T1-weighted surface-reconstruction procedures. In addition, the 7 T structural images showed substantial spatial intensity nonuniformity. We therefore applied two successive intensity-correction steps before constructing the image used for surface placement. First, the original T2-weighted image was corrected for smooth multiplicative bias using the N4 bias-field correction implemented in ANTs [1]. The resulting N4-corrected image served as the structural input for all subsequent reconstruction steps. N4 substantially reduced large-scale intensity inhomogeneity while preserving the native T2-weighted tissue contrast. Residual spatial intensity variation remained after N4 correction, particularly in regions affected by pronounced transmit-field inhomogeneity at 7 T. A second, local normalization was therefore applied before tissue-specific intensity mapping. Let $I_{\mathrm{N4}}(\mathbf{x})$ denote the N4-corrected T2-weighted image and $M$ the brain mask. A smoothly varying local reference intensity was estimated as:

$$r(\mathbf{x}) = \frac{G_\sigma(I_{\mathrm{N4}}\mathbf{1}_M)(\mathbf{x})}{\max\left[G_\sigma(\mathbf{1}_M)(\mathbf{x}), \epsilon\right]}, \tag{S1}$$

where $G_\sigma$ denotes Gaussian smoothing with $\sigma = 8$ mm and $\epsilon$ is a small positive constant preventing numerical instability near the mask boundary. The locally normalized image was then:

$$g(\mathbf{x}) = \frac{I_{\mathrm{N4}}(\mathbf{x})}{r(\mathbf{x})} = I_{\mathrm{N4}}(\mathbf{x}) \frac{\max\left[G_\sigma(\mathbf{1}_M)(\mathbf{x}), \epsilon\right]}{G_\sigma(I_{\mathrm{N4}}\mathbf{1}_M)(\mathbf{x})}, \qquad \mathbf{x} \in M. \tag{S2}$$

Thus, whereas N4 corrected the smooth global bias field, the subsequent Gaussian normalization compensated for residual local intensity variation that remained problematic for tissue-dependent intensity thresholds and surface placement.

Reference WM and GM intensities were estimated from the interior of the corresponding tissue masks,

$$g_{\mathrm{WM}} = \mathrm{median}_{\mathbf{x} \in W^{-2}}\, g(\mathbf{x}), \qquad g_{\mathrm{GM}} = \mathrm{median}_{\mathbf{x} \in G^{-1}}\, g(\mathbf{x}), \tag{S3}$$

where $W^{-2}$ and $G^{-1}$ denote two-step WM and one-step GM erosions, respectively. Erosion reduced contamination from partial-volume voxels near tissue boundaries, and median intensities were used to limit sensitivity to residual local outliers. The locally normalized image was then transformed to the WM-bright/GM-darker ordering expected by conventional T1-weighted surface-reconstruction procedures:

$$T(g) = \begin{cases} 110\, g/g_{\mathrm{WM}}, & g \leq g_{\mathrm{WM}}, \\ 110 - 50\dfrac{g - g_{\mathrm{WM}}}{g_{\mathrm{GM}} - g_{\mathrm{WM}}}, & g > g_{\mathrm{WM}}. \end{cases} \tag{S4}$$

This mapping placed the reference WM and GM intensities near 110 and 60, respectively, while retaining MRI-derived intensity variation around tissue boundaries. To further stabilize white-surface placement, the transformed image was combined with a label-derived reference image $L$, in which WM and GM were assigned intensities of 110 and 60:

$$B_{50} = 0.5\, T(g) + 0.5\, L. \tag{S5}$$

The resulting $B_{50}$ image therefore combined locally normalized MRI contrast with segmentation-derived tissue priors and was used as the intensity target for white-surface deformation.

##### S2.1.3 White-surface reconstruction

An initial WM tessellation was generated from the prepared WM mask on the 0.5-mm working grid. At this resolution, direct tessellation produced a densely sampled mesh containing many short edges and small voxel-scale irregularities. Performing topology correction directly on this mesh increased sensitivity to these local features without providing corresponding anatomical information.

The initial mesh was therefore simplified using quadric error-based edge-collapse decimation [2]. Decimation was continued until the mesh reached a target mean edge length of approximately 1 mm. This reduced representation preserved the overall geometry of the cortical boundary while suppressing small voxel-scale irregularities and reducing the complexity of subsequent topology correction. Topological defects were then identified and corrected on the simplified mesh using the topology-correction framework of Fischl and colleagues [3], yielding a surface with the expected spherical topology for a cerebral hemisphere. After topology correction, the white surface was deformably positioned toward the GM–WM boundary using the $B_{50}$ image as the intensity target [4]. The deformation therefore combined the topologically corrected geometric initialization with the locally varying MRI-derived boundary contrast encoded in $B_{50}$. Following white-surface placement, the topology-corrected mesh was remeshed to an approximately 0.5-mm target edge length. This finer sampling was used for subsequent cortical-depth and local surface analyses. Mesh geometry was regularized during this stage to reduce irregular triangle configurations introduced by tessellation, topology correction, and remeshing while retaining the reconstructed GM–WM boundary.

##### S2.1.4 Recovery of sulcal spaces

Pial-surface reconstruction required an additional preprocessing step because fixation altered the geometry of many narrow sulci. In several locations, opposing cortical banks were brought into close apposition, leaving little or no visible fluid space between them in the acquired volume. Under these conditions, a conventional intensity-driven pial deformation can fail to distinguish the two banks and may bridge across the sulcus. Collapsed or poorly resolved sulcal spaces were therefore recovered in the volumetric domain using the `RecoverSulci` procedure implemented in iBEAT V2.0 [5]. The procedure was applied after reconstruction of the white surface and before pial-surface deformation. Voxels identified as recovered sulcal space were assigned a CSF-like intensity in the image used to guide the pial boundary. This introduced an explicit low-intensity compartment between opposing cortical banks and provided the deformable surface with a volumetric target corresponding to the expected GM–fluid interface. This step was intended to restore separability of closely apposed sulcal banks rather than to alter the position of the reconstructed white surface.

##### S2.1.5 Pial-surface reconstruction

The pial surface was initialized from the topology-corrected white surface and subsequently estimated using the deformable surface model implemented in `mris_place_surface` [4]. The surface was driven outward from the white matter toward the reconstructed GM–fluid boundary, using the sulcal-space-corrected image as the volumetric reference. This deformation differs from a procedure in which each vertex is displaced independently along a fixed white-surface normal. The surface model includes geometric regularization of neighboring vertices and permits both normal and tangential adjustments during deformation. The tangential degrees of freedom are particularly important in narrow and highly curved sulci, where the location of a pial vertex cannot generally be represented by displacement along the corresponding white-surface normal alone. The resulting deformation allowed vertices to move toward sulcal fundi while maintaining a smooth and coherent cortical sheet. The final white and pial surfaces were inspected as paired representations of the GM–WM and GM–fluid boundaries. Together they defined the cortical ribbon used for all subsequent surface-based sampling, including cortical-thickness estimation, cortical-depth profiling, and projection of diffusion-derived measures.

##### S2.1.6 Surface quality control

All reconstructed white and pial surfaces were visually inspected in each specimen before downstream analysis. Surface contours were examined over the anatomical T2-weighted image and the corresponding reconstruction images in multiple orthogonal planes, with particular attention to highly curved gyral crowns, narrow sulci, and regions near the medial cutting plane. Quality control focused on white-surface displacement into cortical GM, pial leakage beyond the cortical boundary, bridging between opposing sulcal banks, and local inconsistencies between the reconstructed surfaces and the underlying tissue segmentation. All six reconstructed hemispheres passed the final visual quality-control assessment and were included in subsequent cortical-thickness estimation, cortical-depth sampling, and diffusion analyses.

#### S2.2 Gyral-skeleton construction and landmark extraction

Three-hinge gyral landmarks were extracted from the reconstructed white surface using the automated gyral-skeleton framework described previously [6]. Cortical folding patterns were first characterized using vertex-wise sulcal depth and mean curvature. Sulcal depth values were used to distinguish gyral and sulcal regions, with positive values indicating gyri and negative values indicating sulci. To improve robustness, isolated vertices were removed and minor inconsistencies in sulcal classification were corrected using local curvature information. A gyral thinning procedure was then applied to extract the central ridges of gyri. Specifically, gyral regions were iteratively thinned from sulcal boundaries under curvature constraints, progressively reducing each gyral region to a narrow ridge representing the gyral skeleton. This skeleton approximates the medial axis of gyral structures and captures the topological organization of cortical folding patterns. A skeleton graph was subsequently constructed by connecting neighboring vertices along the gyral skeleton. The graph was further refined by removing short branches and spurious connections to preserve only meaningful gyral ridges. In addition, long gyral ridges that were not initially connected to the skeleton were detected using curvature-based criteria and linked to the skeleton through shortest paths on the cortical surface. Finally, 3HGs were identified as vertices on the gyral skeleton with three branches, corresponding to junctions where multiple gyral ridges converge. To ensure topological consistency, nearby hinge pairs were further refined using surface-constrained shortest-path rewiring to eliminate redundant hinge detections.

##### S2.3 Tractography reconstruction

Fiber orientation distributions were independently estimated from the preprocessed HARDI data using DIPY [7]. A single-shell single-tissue response function was estimated using `auto_response_ssst`, followed by constrained spherical deconvolution with maximum spherical-harmonic order $l_{\max} = 8$. Fiber-orientation peaks were extracted on a 724-direction sphere. Peaks were retained when their amplitude exceeded 50% of the largest local peak and when neighboring peaks were separated by at least 25°; up to three peaks were retained per voxel. Deterministic streamline propagation used DIPY `LocalTracking` with the resulting `PeaksAndMetrics` representation as an EuDX-based direction getter. Seeds were placed throughout the white-matter mask at one seed per voxel. In the primary reconstruction, at most one admissible initial direction was propagated from each seed (`max_cross`=1). Tracking used a 0.4-mm step size and a maximum turning angle of 60°. A diffusion-tensor model was fitted separately to derive FA for streamline termination only; tensor eigenvectors were not used as tracking directions. Tracking stopped when FA fell below 0.12 or when no admissible propagation direction remained. Streamline fragments containing fewer than six tracking points were discarded.

##### S2.4 Connectivity node definition and endpoint assignment

Connectivity nodes were defined in superficial white matter rather than directly on the cortical surface. For each gyral junction, a spherical patch of 3-mm radius was centered 2 mm beneath the white surface along the local surface normal. Junctions separated by less than 6 mm were merged and represented by the cluster medoid. Each retained 3HG node was paired with one gyral-crown control from the same DK parcel. Crown nodes were required to remain spatially separated so that the two node classes had comparable node resolution. Junctions without an eligible within-parcel crown control were omitted from the paired connectivity analysis. A streamline was assigned to a seed when it intersected the 3-mm seed neighborhood. The endpoint distal to the seed was then identified. A streamline was considered to reach a target node when this distal endpoint lay within 4 mm of the target-patch center. For seed $i$ and target $j$, connectivity was defined as:

$$F_{ij} = \frac{N_{ij}}{N_i}, \tag{S6}$$

where $N_i$ denotes the number of streamlines sampled from seed $i$ and $N_{ij}$ denotes the subset whose distal endpoint reached target $j$.

#### S2.5 Partial-volume sensitivity analysis

Although the diffusion acquisition provided substantially higher spatial resolution than conventional in vivo diffusion MRI, its voxel size remained larger than that of the structural image and reconstructed cortical geometry. We therefore tested whether the intracortical 3HG–crown differences could arise from systematic differences in GM/WM partial volume at tightly folded gyral junctions. A three-class tissue volume was constructed at the 0.5-mm structural resolution using the reconstructed white surface and cortical ribbon. Voxels inside the white surface were assigned to WM, voxels within the cortical ribbon to cortical GM, and remaining voxels to a third non-GM/non-WM class. Rather than interpolating the categorical tissue masks to diffusion resolution, every structural voxel was assigned to the diffusion voxel containing its center. Tissue-class counts were then accumulated within each diffusion voxel, yielding exact volumetric fractions of GM, WM, and remaining tissue. Each equivolumetric cortical sample was associated with the tissue fractions of the diffusion voxel from which its MRI value was obtained. The region-matched cortical-depth analysis was then repeated under three conditions.

First, the original analysis used all samples irrespective of tissue composition. Second, a high-purity analysis retained only samples obtained from diffusion voxels containing at least 70% cortical GM. Third, an explicit partial-volume adjustment was performed independently within each DK parcel and cortical depth using

$$Y = \beta_0 + \beta_1\,\mathrm{class} + \beta_2\, f_{\mathrm{WM}} + \beta_3\, f_{\mathrm{other}} + \epsilon, \tag{S7}$$

where $\mathrm{class}$ indicated 3HG versus crown and $f_{\mathrm{WM}}$ and $f_{\mathrm{other}}$ were the corresponding tissue fractions. The adjusted class coefficient $\beta_1$ was retained as the 3HG–crown contrast. The comparison of the original, high-GM-purity, and covariate-adjusted estimates was used specifically as a sensitivity analysis; the primary depth-resolved statistics remained those defined in the main Methods.

#### S2.6 Control for gyral tractography bias

Diffusion tractography is known to exhibit a gyral termination bias, whereby streamline endpoints preferentially accumulate near gyral crowns. Because 3HGs and their crown controls occupy different positions within the gyral folding profile, we quantified this bias directly and examined how strongly it propagated through the sub-WM connectivity analysis. Each streamline endpoint was assigned to its nearest white-surface vertex when the endpoint lay within 3 mm of the surface. Endpoint counts were accumulated across 20 percentile bins of cortical sulcal depth. To account for unequal amounts of cortical surface represented by different folding positions, the number of endpoints in each bin was divided by its cortical surface area,

$$D_k = \frac{N_k^{\mathrm{endpoint}}}{A_k}, \tag{S8}$$

where $D_k$ is endpoint density in sulcal-depth bin $k$, $N_k^{\mathrm{endpoint}}$ is the number of assigned streamline endpoints, and $A_k$ is the corresponding cortical surface area. The median folding positions of the 3HG and matched crown node classes were located on this endpoint-density profile. We then quantified propagation of the class difference through three stages of the connectivity analysis: cortical endpoint density, number of streamlines collected by each 3-mm sub-WM patch, and graph node degree. To assess the effect on pairwise connection incidence, a logistic regression was fitted separately within each specimen. For node pair $(i, j)$,

$$\mathrm{logit}\, P(A_{ij} = 1) = \beta_0 + \beta_1\, C_{ij} + f(D_{ij}) + \beta_2\, S_{ij} + \beta_3 \log N_i, \tag{S9}$$

where $A_{ij}$ indicates whether a connection was detected, $C_{ij}$ represents node class, $D_{ij}$ is inter-node distance, $S_{ij}$ represents folding position/sulcal depth, and $N_i$ is the number of streamlines sampled by the seed node. The class effect was compared with models omitting the streamline-count term.

This analysis was used to determine which connectivity estimator was least sensitive to sampling density. Because the raw pair-incidence effect was strongly attenuated after adjustment for the number of streamlines sampled at each node, pair incidence was not used as the primary evidence for preferential inter-hinge wiring. Instead, the primary analysis used the seed contrast:

$$E_{\text{seed}} = \frac{f_{HH}}{f_{CH}}, \tag{S10}$$

for which both classes are compared against the same 3HG target set and each seed is normalized by its own sampled streamline count.

#### S2.7 Tractography parameter sensitivity

We evaluated whether the preferential inter-hinge connectivity pattern depended on the particular parameterization used for deterministic tractography. The stored CSD orientation peaks were first reconstructed within the tracking framework. Because each stored peak direction corresponded to a vertex on the sampling sphere, its original sphere index was recovered by maximum absolute directional correspondence. The reconstructed direction getter was verified against the reference tracking configuration before any parameter was varied. Using the reference settings reproduced the stored tractogram streamline for each streamline. The reference configuration used a maximum turning angle of $60^\circ$, a 0.4-mm tracking step, a peak-weight parameter of 0.50, and an FA stopping threshold of 0.12. Starting from this configuration, tracking was repeated while independently varying six aspects of the procedure:

- the FOD cutoff;
- the tracking peak-weight threshold;
- the maximum turning angle;
- the tracking step size;
- the FA stopping threshold; and
- the number of seeds.

The tested alternatives included FOD cutoffs of 0.65 and 0.80, peak-weight thresholds of 0.30 and 0.70, turning angles of $45^\circ$ and $75^\circ$, a step size of 0.6 mm, an FA stopping threshold of 0.15, and an eightfold increase in seeding density. FOD cutoffs below the peak-extraction threshold of 0.5 were not evaluated because doing so would have required refitting the original CSD peak representation. For every tracking configuration, the complete node construction and connectivity analysis were repeated using the same 3HG junctions, same-parcel crown controls, 3-mm sub-WM patches, and 4-mm target-endpoint tolerance as in the primary analysis. The target contrast:

$$E_{\text{target}} = \frac{f_{HH}}{f_{HC}}, \tag{S11}$$

and the seed-normalized contrast:

$$E_{\text{seed}} = \frac{f_{HH}}{f_{CH}}, \tag{S12}$$

were recomputed independently for each specimen and parameter configuration. This analysis tested whether the direction and magnitude of preferential inter-hinge wiring were preserved across reasonable changes in the tractography parameters rather than being specific to a single tracking configuration.

##### S2.8 Tractography reconstruction sensitivity

We additionally tested whether the inter-hinge connectivity preference depended on assumptions specific to the reference streamline reconstruction. First, we examined the number of initial directions propagated from each white-matter seed. The reference reconstruction used `max_cross`=1, such that at most one initial FOD peak was followed from each seed. Because multiple local fiber orientations may occur near gyral junctions, tracking was repeated with `max_cross`=2, `max_cross`=3, and with all admissible initial directions. All other tracking parameters and the stored CSD peaks were held fixed. The reconstructed EuDX direction getter was first verified to reproduce the reference tractogram streamline for each streamline before `max_cross` was varied. Second, we tested sensitivity to the rule used to select the local propagation direction. For each specimen, a common CSD FOD representation was used to generate three tractograms with identical white-matter seeds, an FA stopping threshold of 0.12, a maximum turning angle of 60°, and a step size of 0.4 mm. The three reconstructions differed only in whether propagation followed discrete FOD peaks, the local maximum of the continuous FOD, or probabilistic sampling from the FOD. To assess dependence on software implementation, tractography was also reconstructed independently in MRtrix3 [8] using `dwi2response tournier`, `dwi2fod csd` ($l_{\max} = 8$), and deterministic `SD_Stream` tracking seeded from the same white-matter mask. The FOD amplitude cutoff was fixed at 0.05 before connectivity contrasts were computed. All tractograms were evaluated using the same node geometry and seed-normalized connectivity estimator as in the primary analysis. Specifically, the contrast quantified the fraction of streamlines sampled by a seed node whose distal endpoint fell within 4 mm of a 3HG target node. Specimen-level seed contrasts were log-transformed and evaluated across the six specimens using two-sided one-sample $t$-tests. MRtrix3 iFOD2 was also evaluated as a probabilistic reconstruction. It was not included in the cross-reconstruction connectivity comparison because its endpoint distribution was incompatible with the endpoint-based node definition used in the present analysis. Without anatomical constraints, the iFOD2 endpoint distribution extended substantially beyond the gray–white matter interface. Its endpoint-depth distribution is nevertheless shown in Supplementary Fig. S3c to make this exclusion explicit.

#### References


[1] Nicholas J Tustison, Brian B Avants, Philip A Cook, Yuanjie Zheng, Alexander Egan, Paul A Yushkevich, and James C Gee. N4itk: improved n3 bias correction. *IEEE transactions on medical imaging*, 29 (6):1310–1320, 2010. doi: 10.1109/TMI.2010.2046908.

[2] Michael Garland and Paul S Heckbert. Surface simplification using quadric error metrics. In *Proceedings of the 24th annual conference on Computer graphics and interactive techniques*, pages 209–216, 1997. doi: 10.1145/258734.258849.

[3] Bruce Fischl, Arthur Liu, and Anders M Dale. Automated manifold surgery: constructing geometrically accurate and topologically correct models of the human cerebral cortex. *IEEE transactions on medical imaging*, 20(1):70–80, 2001. doi: 10.1109/42.906426.

[4] Anders M Dale, Bruce Fischl, and Martin I Sereno. Cortical surface-based analysis: I. segmentation and surface reconstruction. *Neuroimage*, 9(2):179–194, 1999. doi: 10.1006/nimg.1998.0395.

[5] Li Wang, Zhengwang Wu, Liangjun Chen, Yue Sun, Weili Lin, and Gang Li. ibeat v2. 0: a multisite-applicable, deep learning-based pipeline for infant cerebral cortical surface reconstruction. *Nature protocols*, 18(5):1488–1509, 2023. doi: 10.1038/s41596-023-00806-x.

[6] Minheng Chen, Chao Cao, Tong Chen, Dunia Alhamad, Tianming Liu, Li Su, and Dajiang Zhu. Fine-scale individualized gyral folding-based cortical similarity networks reveal distinct organizational pat-

terns in alzheimer's disease and lewy body dementia. *Imaging Neuroscience*, 4:IMAG.a.1322, 08 2026. ISSN 2837-6056. doi: 10.1162/IMAG.a.1322.

[7] Eleftherios Garyfallidis, Matthew Brett, Bagrat Amirbekian, Ariel Rokem, Stefan Van Der Walt, Maxime Descoteaux, and Ian Nimmo-Smith. Dipy, a library for the analysis of diffusion mri data. *Frontiers in Neuroinformatics*, Volume 8 - 2014, 2014. ISSN 1662-5196. doi: 10.3389/fninf.2014.00008.

[8] J-Donald Tournier, Robert Smith, David Raffelt, Rami Tabbara, Thijs Dhollander, Maximilian Pietsch, Daan Christiaens, Ben Jeurissen, Chun-Hung Yeh, and Alan Connelly. Mrtrix3: A fast, flexible and open software framework for medical image processing and visualisation. *Neuroimage*, 202:116137, 2019. doi: 10.1016/j.neuroimage.2019.116137.
